# Ether-lipid deficiency remodels the testicular interstitium beyond spermatogenic arrest in *Gnpat*-knockout mice

**DOI:** 10.64898/2026.10.17.759782

**Authors:** Niwase Shamim, Akihisha Fujiyama, Masanori Honsho, Shiro Mawatari, Takehiko Fujino

## Abstract

Ether lipids, including plasmalogens and the germ-cell glycolipid seminolipid, require the peroxisomal enzyme GNPAT. *Gnpat*-knockout mice are male-infertile, but their testis has been described only qualitatively. We immunostained testis sections from 12-week-old wild-type (WT) and *Gnpat*-knockout (KO) mice (n = 3 per genotype) for SOX9 and *γ*H2AX with DAPI and analysed 24 fields (22,560 nuclei) with an automated pipeline and a conservative animal-level analysis. Seminiferous tubules fell from 90% to 43% of the section, and the interstitium expanded five-fold (from 11% to 54%). Interstitial cells were 1.6-fold larger but not more densely packed, and half of them, against a quarter in WT, showed lipofuscin-like autofluorescence. Spermatid-sized nuclei fell from 38% to 10%, with the loss confined to the adluminal compartment. The architectural, interstitial and germ-cell changes held at the animal level (*P* ≤ 0.02). Anucleate SOX9-immunoreactive bodies, likely residual bodies, were 2.2-fold rarer, and *γ*H2AX was not increased, which argues against a DNA-damage checkpoint arrest. Ether-lipid deficiency thus arrests germ cells, most likely through loss of seminolipid, and remodels the interstitium into an expanded compartment of enlarged, autofluorescent cells, a second target relevant to rhizomelic chondrodysplasia punctata.

## Introduction

Ether lipids carry a fatty alcohol linked by an ether bond at the *sn*-1 position of glycerol. Their largest class, the plasmalogens, make up about one-fifth of phospholipid mass in humans and shape membrane curvature, lipid rafts, antioxidant defence and cholesterol homeostasis [1–4]. Their synthesis begins in the peroxisome, where glyceronephosphate O-acyltransferase (GNPAT) acylates dihydroxyacetone phosphate and alkylglycerone-phosphate synthase (AGPS) introduces the ether bond with a fatty alcohol supplied by fatty-acyl-CoA reductase 1 (FAR1); the backbone is completed in the endoplasmic reticulum, where PEDS1 creates the vinyl-ether bond of plasmalogens (Fig. 1a) [1,5,6]. Defects in GNPAT, AGPS or the import receptor PEX7 cause rhizomelic chondrodysplasia punctata (RCDP), a severe multisystem disorder [1,7].

**Figure 1.**
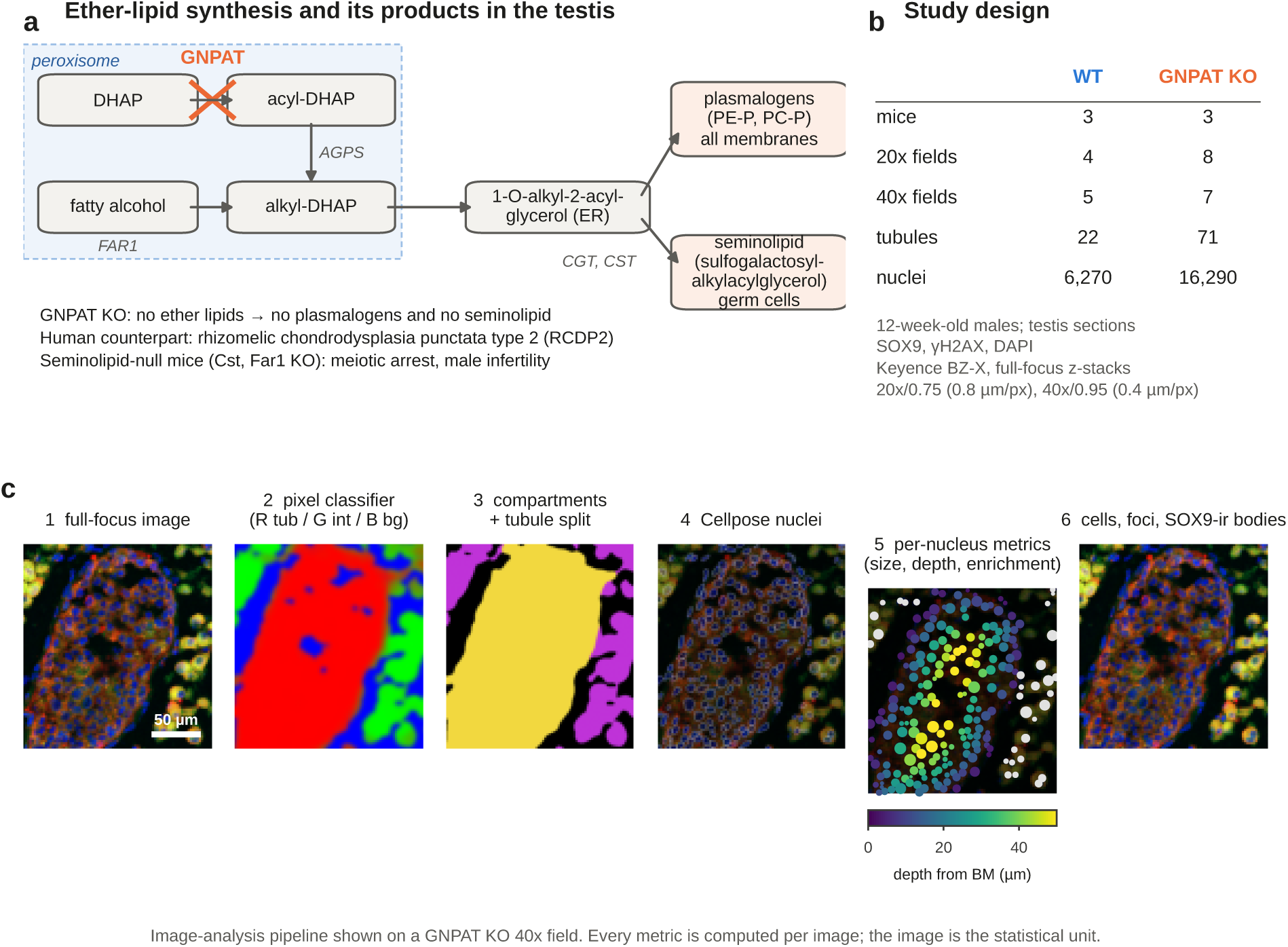
Background, study design and image-analysis pipeline. **(a)** Ether-lipid synthesis and its products in the testis. In the peroxisome, GNPAT acylates dihydroxyacetone phosphate (DHAP) to acyl-DHAP. AGPS exchanges the acyl chain for a fatty alcohol supplied by FAR1, forming alkyl-DHAP. The resulting 1-*O*-alkyl-2-acyl-glycerol is completed in the endoplasmic reticulum (ER) to plasmalogens (PE-P, PC-P), present in all membranes, or, through CGT and CST, to seminolipid (sulfogalactosyl-alkylacylglycerol) in germ cells. GNPAT loss therefore abolishes both. **(b)** Study design: 3 mice per genotype, number of 20× and 40× fields, valid tubule profiles and segmented nuclei; acquisition settings below. **(c)** Pipeline applied to a *Gnpat*-KO 40× field: (1) extended-focus image, red (SOX9); green (*γ*H2AX); blue (DAPI); (2) random-forest probability map (red, tubule; green, interstitium; blue, background); (3) compartments after tubule splitting (yellow, tubule profiles; magenta, interstitium); (4) Cellpose nuclei (outlines); (5) per-nucleus metrics (tubular nuclei coloured by depth from the basement membrane, BM, scale bar below; grey, interstitial nuclei; marker size proportional to nuclear area); (6) interstitial cell territories, *γ*H2AX foci and SOX9-immunoreactive (SOX9-ir) bodies (cyan). Scale bar, 50 µm. Every metric is computed per image, and the image (field) is the statistical unit.

The testis is a revealing tissue in which to study ether lipids. Developing germ cells make seminolipid (sulfogalactosyl-alkylacylglycerol), which accounts for more than 90% of testicular glycolipid and whose alkyl chain is built by the same GNPAT–AGPS–FAR1 pathway [8–10]. Mice that cannot galactosylate or sulfate it (*Cgt*- or *Cst*-null), or that lack *Far1*, arrest at late pachytene or metaphase I [8,10,11]. *Gnpat*-null males are infertile, with atrophic tubules, apoptotic pachytene spermatocytes, multinucleated giant cells and no elongated spermatids; the original report also noted “proliferating Leydig cell clusters” [12,13], and ether-lipid deficiency disrupts remodelling of the blood–testis barrier [14]. *Agps*-deficient and *Pex7*-null mice are likewise male-sterile [15,16], and germ-cell-specific loss of peroxisomes arrests spermatogenesis [17,18].

Three gaps remain. Every account of the *Gnpat*-null testis is qualitative, so the size and variability of the phenotype, and especially the interstitial changes, have never been measured. Fluorescence imaging has not been applied, so properties such as autofluorescent pigment have not been examined. And which ether lipid matters is unresolved: mice lacking only plasmalogens (*Peds1*-null) are fertile [6], whereas ether-lipid-deficient cells mishandle cholesterol [19,20], a concern for the steroidogenic interstitium. Here we quantify the *Gnpat*-knockout testis with SOX9/*γ*H2AX/DAPI immunofluorescence and an automated image-analysis pipeline. We confirm the spermatogenic phenotype and report three new findings: a five-fold expansion of the interstitium made up of enlarged cells, a large population of autofluorescent interstitial cells, and loss of anucleate SOX9-immunoreactive round bodies.

## Results

### An automated pipeline for whole-field testis histomorphometry

Because GNPAT catalyses the first step of ether-lipid synthesis, its loss removes both plasmalogens and seminolipid (Fig. 1a). We immunostained testis cryosections from three WT and three *Gnpat*-KO male mice at 12 weeks of age, the age examined in the original description of *Gnpat*-deficient mice [12], for SOX9 and *γ*H2AX with a DAPI counterstain, and imaged 24 extended-focus fields (WT, 4 at 20× and 5 at 40×; KO, 8 at 20× and 7 at 40×; Fig. 1b). Every field was processed by the same automated pipeline (Fig. 1c; Methods). A random-forest pixel classifier separated seminiferous tubules, interstitium and background, a distance-transform watershed split touching tubule profiles, and Cellpose segmented every nucleus; each nucleus was then assigned a compartment, a depth from the basement membrane (BM) and a marker content, and interstitial cell territories, *γ*H2AX foci and SOX9-immunoreactive bodies were detected. The fields contained 22,560 segmented nuclei (6,270 WT and 16,290 KO) and 93 valid tubule profiles (22 WT and 71 KO). Segmentation overlays confirmed that tubule outlines followed the BM and that the broad KO interstitium was captured completely (Fig. S1). The *γ*H2AX-channel exposure was identical between genotypes at 20× (150 ms) but not at 40× (170 ms WT, 350 ms KO; Fig. S2a–c), so raw intensities were never compared between genotypes.

### The *Gnpat*-null testis is dominated by interstitium

The genotypes differed at a glance (Fig. 2). At 20×, WT sections were filled with large, closely apposed tubules whose multilayered epithelium contained many small spermatid nuclei, separated by narrow interstitial strips (Fig. 2a). KO sections instead contained small, round tubules separated by broad sheets of large interstitial cells (Fig. 2b). At 40×, the WT epithelium contained many compact, intensely SOX9-immunoreactive round bodies (Fig. 2c), whereas the KO interstitium consisted of rounded to polygonal cells with abundant granular cytoplasm (Fig. 2d). Single-channel crops of identical size and scale (120 × 120 µm) from matched compartments showed diffuse cytoplasmic SOX9-channel signal with compact bodies and a weak, diffuse *γ*H2AX signal within WT tubules (Fig. 2e). KO interstitial cells, by contrast, were bright in both the SOX9 and the *γ*H2AX channel, with the same cytoplasmic distribution in each (Fig. 2f). Two different antibodies would not label the same cytoplasm with matched intensity through two filter sets, so this pattern is the hallmark of broad-spectrum autofluorescence, which we quantify below.

**Figure 2.**
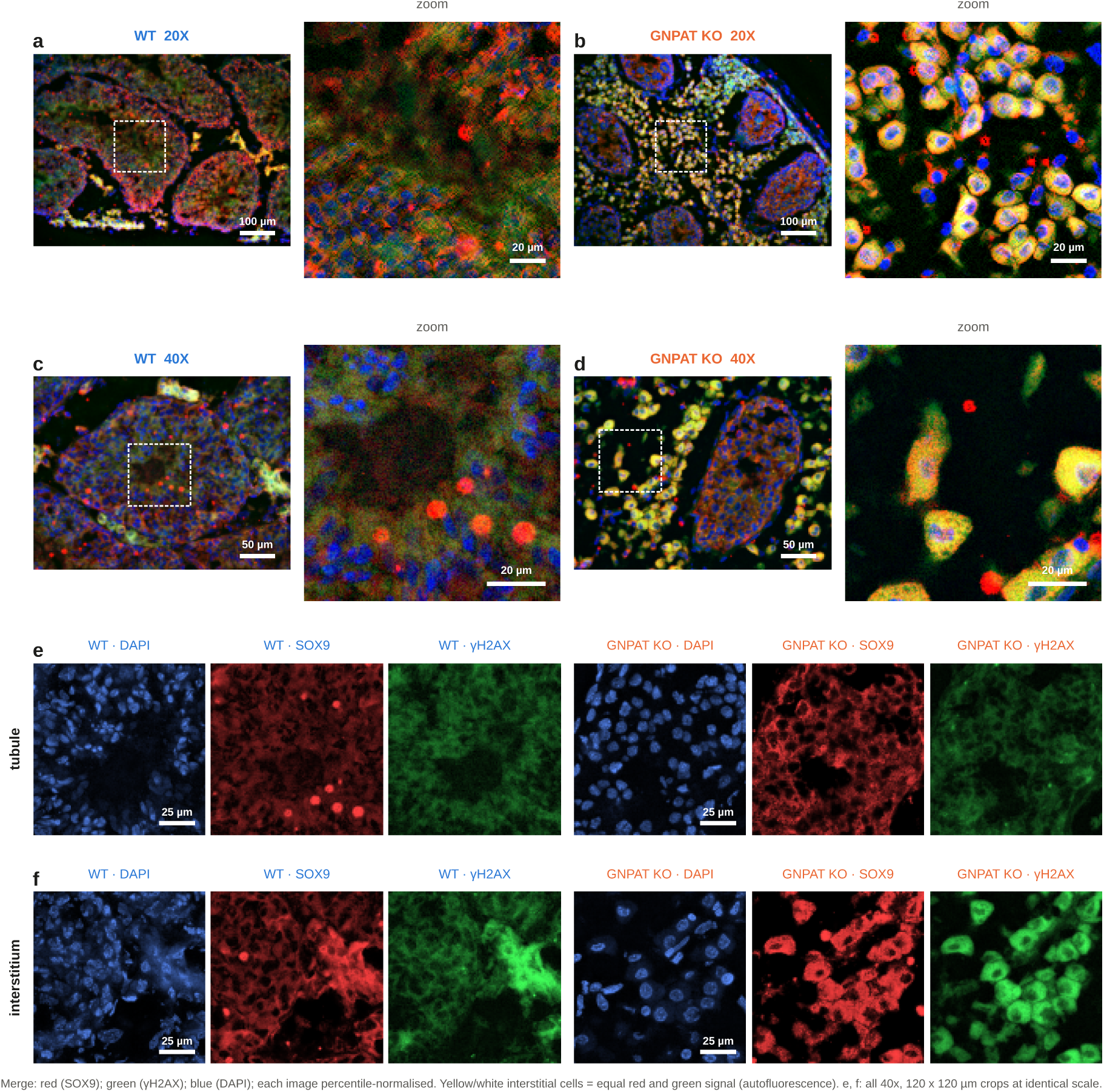
Representative immunofluorescence of WT and *Gnpat*-KO testis. Merged images: red (SOX9); green (*γ*H2AX); blue (DAPI); each image percentile-normalised. **(a, b)** 20× fields of WT (a) and KO (b) testis, each with an enlarged view (right) of the region in the dashed box. WT: large, closely apposed tubules with a multilayered epithelium and narrow interstitium. KO: small round tubules separated by broad sheets of large interstitial cells that appear yellow to orange. **(c, d)** 40× fields of WT (c) and KO (d) with enlarged views. In WT, compact SOX9-immunoreactive round bodies lie within the epithelium. In KO, the interstitium is formed by rounded to polygonal cells with granular cytoplasm. **(e, f)** Single channels of the 40× fields shown in (c, d), in matched compartments: blue (DAPI), red (SOX9), green (*γ*H2AX). **(e)** The most tubule-rich 120 × 120 µm window of each field. Nuclear size and density are comparable between genotypes; WT shows compact SOX9-immunoreactive bodies. **(f)** The most interstitium-rich window. KO interstitial cells are equally bright in the SOX9 and *γ*H2AX channels (yellow to white in the merge), indicating broad-spectrum autofluorescence rather than antibody labelling. All crops in (e, f) are at identical scale and were taken with the same 40×/0.95 objective without digital zoom. Scale bars: 100 µm (20× fields), 50 µm (40× fields), 20 µm (enlarged views), 25 µm (e, f).

### Tubular atrophy

Tubules occupied a median 89.6% of the WT section but only 42.8% of the KO section (Fig. 3a, b; difference −46.5 percentage points, 95% CI −52.4 to −40.6, *P*_adj_ < 0.001), and the interstitium expanded reciprocally from 11.1% to 53.9% (Fig. 3c; +42.0 points, 95% CI 36.3 to 47.7, *P*_adj_ < 0.001). Field-level values did not overlap between genotypes, and both effects held in the conservative animal-level analysis (*P*_animal_ = 0.001; Table 1). KO tubules were narrower, with a median inscribed diameter of 104 vs. 149 µm per field (Fig. 3d; *P*_adj_ = 0.007); a tubule-level mixed model across all 93 profiles gave a difference of −51 µm (95% CI −76 to −26), and the whole distribution of individual diameters was shifted to smaller values (Fig. 3e). This reduction of about 30% is a lower bound, because every WT profile but only 69% of KO profiles were cut by the field edge (Fig. 3f, squares). KO profiles were also more circular (median 0.50 vs. 0.41; Fig. 3f, g; *P*_adj_ = 0.003) and more numerous per tissue area (26 vs. 15 per mm²; Fig. 3h), as expected when the same area contains smaller tubules.

**Figure 3.**
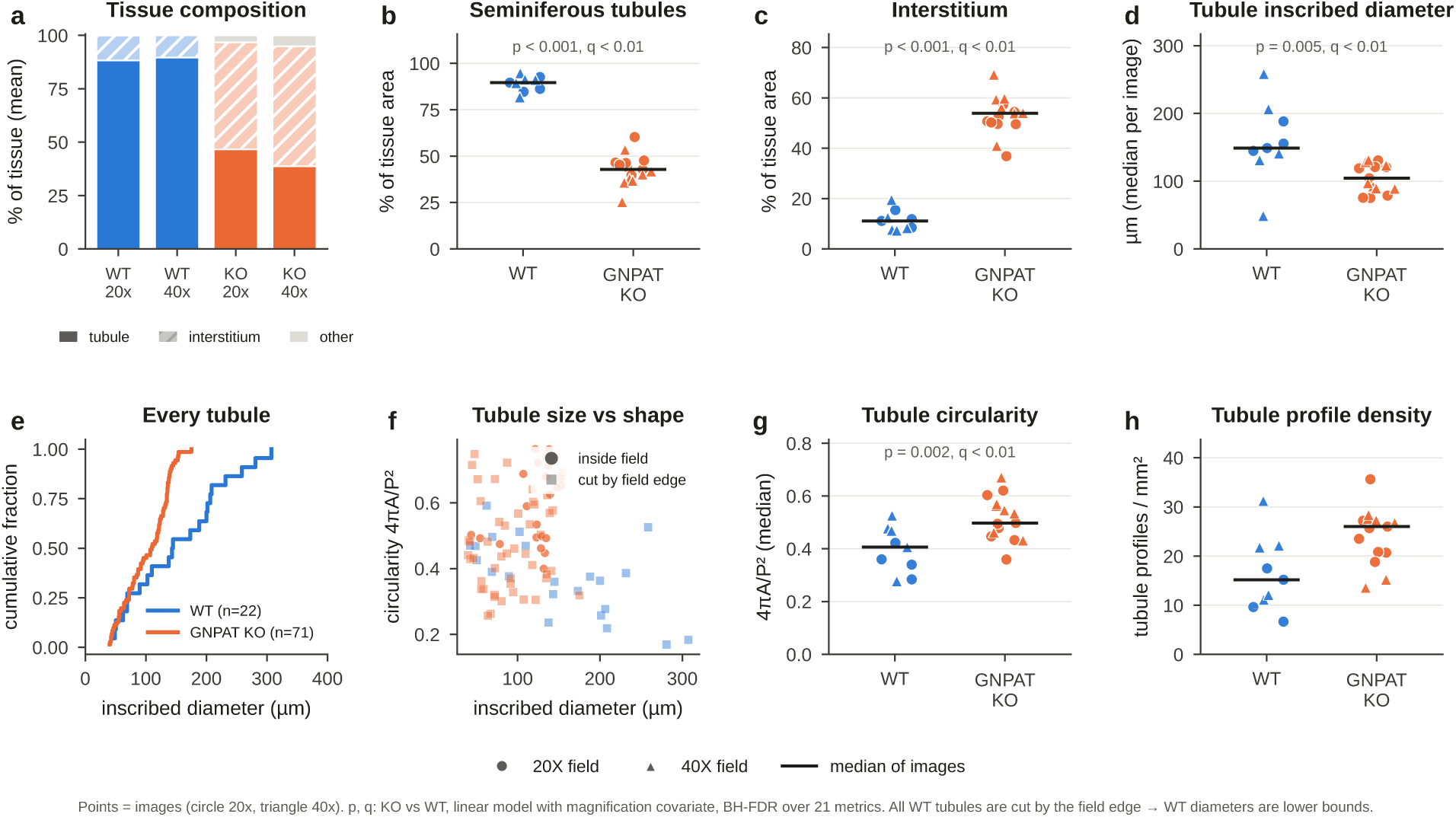
Tubular atrophy and interstitial expansion. **(a)** Mean tissue composition of each field as % of tissue (tubule, interstitium, other), per genotype and magnification. **(b, c)** Seminiferous tubule area (b) and interstitium area (c) as % of tissue per field. **(d)** Median inscribed diameter of valid tubule profiles per field. **(e)** Cumulative distribution of the inscribed diameter of every valid tubule profile (WT n = 22, KO n = 71). **(f)** Tubule size (inscribed diameter) vs. shape (circularity, 4π·area/perimeter²) for each profile. Circles, profiles inside the field; squares, profiles cut by the field edge (all WT profiles), whose diameters are lower bounds. **(g)** Median tubule circularity per field. **(h)** Tubule profiles per mm² of tissue. In (b–d, g, h), each symbol is one field (circles, 20×; triangles, 40×), and bars show the median of fields. Values printed in the panels: *p*, field-level linear model with magnification as covariate; *q*, Benjamini– Hochberg-adjusted *P* value (*P*_adj_) over 21 metrics. Animal-level statistics are given in Table 1.

**Table 1.** Field-level and animal-level comparison of WT and *Gnpat*-KO testis. Values are medians of fields. Δ (95% CI) is from a linear model with magnification as covariate. *P*_adj_ is the Benjamini–Hochberg FDR over 21 metrics. “Animal-level *P*” gives the median and the largest (*P*_animal_) two-sample *t*-test *P* value over all 35.3 million balanced groupings of fields into 3 + 3 mice; * marks *P*_animal_ < 0.05. n.a. = not computable (20×-only metric, 4 WT fields). ^†^ Derived metric added after the literature comparison; unadjusted *P*, not part of the 21-metric FDR set.

| Metric | WT | KO | $\Delta$ KO – WT (95% CI) | $P_{\text{adj}}$ | Animal-level $P$<br>(median / $P_{\text{animal}}$) |
| --- | --- | --- | --- | --- | --- |
| Tubule area (% section) | 89.6 | 42.8 | –46.5 (–52.4, –40.6) | < 0.001 | < 0.001 / 0.001 * |
| Interstitial area (% section) | 11.1 | 53.9 | 42.0 (36.3, 47.7) | < 0.001 | < 0.001 / 0.001 * |
| Tubule inscribed diameter ($\mu\text{m}$) | 149 | 104 | –52.9 (–87.6, –18.2) | 0.007 | 0.048 / 0.20 |
| Tubule circularity | 0.41 | 0.50 | 0.12 (0.05, 0.19) | 0.003 | 0.021 / 0.19 |
| Tubular nuclei per $10^4 \mu\text{m}^2$ | 40.6 | 69.2 | 27.9 (17.1, 38.7) | < 0.001 | 0.008 / 0.092 |
| Tubular nuclear area ($\mu\text{m}^2$) | 23.0 | 40.0 | 16.6 (13.6, 19.6) | < 0.001 | < 0.001 / 0.003 * |
| Spermatid-sized nuclei (%) | 38.1 | 10.0 | –27.8 (–34.7, –20.9) | < 0.001 | 0.002 / 0.020 * |
| Epithelial depth, 90th pct ($\mu\text{m}$) | 52.9 | 41.2 | –17.4 (–25.8, –9.1) | < 0.001 | 0.022 / 0.15 |
| Interstitial nuclei per $\text{mm}^2$ | 567 | 2554 | 2039 (1721, 2357) | < 0.001 | < 0.001 / 0.003 * |
| Interstitial nuclei per $\text{mm}^2$ interstitium$^{\dagger}$ | 5633 | 4948 | –627 (–1226, –28) | 0.041$^{\dagger}$ | 0.11 / 0.38 |
| Interstitial cell territory ($\mu\text{m}^2$) | 90 | 147 | 50.9 (37.5, 64.3) | < 0.001 | 0.001 / 0.017 * |
| Autofluorescent interstitial cells (%; 20×) | 25.2 | 50.1 | 24.5 (16.3, 32.8) | < 0.001 | n.a. |
| $\gamma\text{H2AX}$ enrichment, tubule ($\log_2$) | –0.31 | –0.38 | –0.12 (–0.20, –0.04) | 0.008 | 0.017 / 0.22 |
| $\gamma\text{H2AX}$-enriched tubular nuclei (%) | 1.7 | 3.7 | 1.6 (0.4, 2.8) | 0.013 | 0.021 / 0.23 |
| $\gamma\text{H2AX}$ foci per tubular nucleus | 0.08 | 0.04 | –0.12 (–0.42, 0.19) | 0.44 | 0.54 / 0.63 |
| SOX9 enrichment, tubule ($\log_2$) | –1.09 | –1.41 | –0.26 (–0.54, 0.02) | 0.070 | 0.077 / 0.34 |
| Basal SOX9-nuclear cells per $10^4 \mu\text{m}^2$ | 0.80 | 1.90 | 1.7 (0.5, 2.8) | 0.008 | 0.012 / 0.16 |
| SOX9-immunoreactive bodies per $\text{mm}^2$ | 292 | 125 | –141 (–191, –91) | < 0.001 | 0.007 / 0.077 |

### Germ-cell loss is confined to the adluminal compartment

Tubular nuclear size provided a quantitative readout of germ-cell composition (Fig. 4). At 40×, the WT size distribution peaked at 15–20 µm², the size of spermatid nuclei; this peak was absent in KO, whose distribution had a broad maximum at 30–50 µm² (Fig. 4a). Every KO field (7–17%) had fewer spermatid-sized nuclei (< 20 µm²) than any WT field (21–63%), and large nuclei (> 40 µm²) made up most of the KO epithelium (Fig. 4b). The median proportion of spermatid-sized nuclei fell from 38.1% to 10.0% (Fig. 4c; −27.8 points, 95% CI −34.7 to −20.9, *P*_adj_ < 0.001, *P*_animal_ = 0.020), and median tubular nuclear area rose from 23.0 to 40.0 µm² (Fig. 4d; *P*_adj_ < 0.001, *P*_animal_ = 0.003), a 1.66-fold increase (95% CI 1.51–1.84) in a nucleus-level mixed model of 14,219 nuclei. The KO epithelium was thinner, with a 90th percentile of nuclear depth from the BM of 41.2 vs. 52.9 µm (Fig. 4e; *P*_adj_ < 0.001), and more densely nucleated (69.2 vs. 40.6 nuclei per 10^4^ µm²; Fig. 4f; *P*_adj_ < 0.001), consistent with a collapsed epithelium without a lumen.

**Figure 4.**
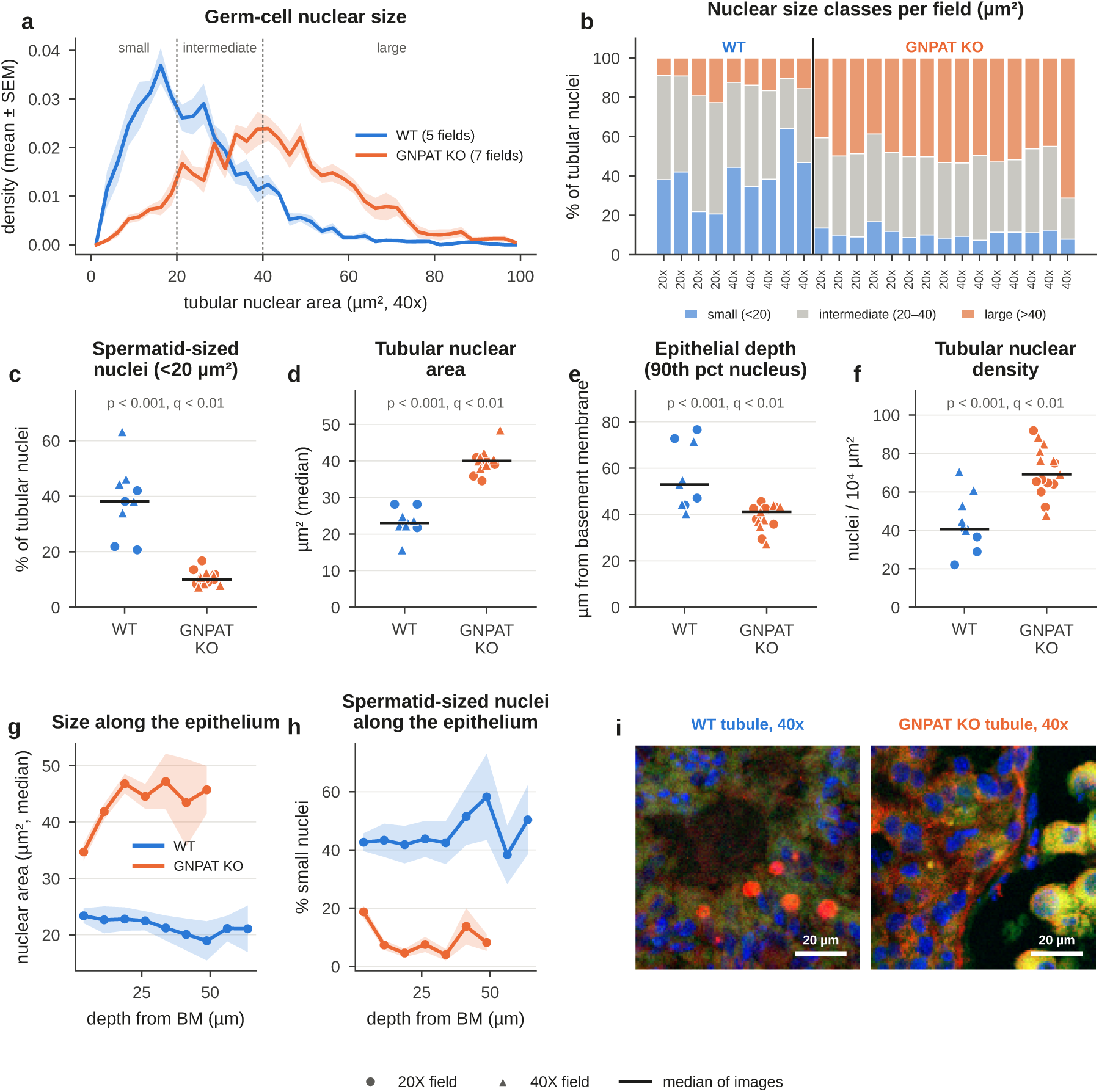
Germ-cell loss is confined to the adluminal compartment. **(a)** Size distribution of tubular nuclei in 40× fields (mean ± SEM of fields; WT 5, KO 7). Dotted lines mark the class boundaries at 20 and 40 µm²; WT shows a spermatid peak at 15–20 µm² that is absent in KO. **(b)** Proportion of small (< 20 µm², spermatid-sized), intermediate (20–40 µm²) and large (> 40 µm²) tubular nuclei in each field (labels give magnification). **(c–f)** Field-level proportion of spermatid-sized nuclei (c), median tubular nuclear area (d), epithelial depth (90th percentile of nuclear distance from the BM) (e) and tubular nuclear density per 10^4^ µm² of tubule (f). Symbols as in Fig. 3 (circles, 20×; triangles, 40×; bar, median). **(g, h)** Radial profiles from the BM towards the lumen in 40× fields: median nuclear area (g) and proportion of spermatid-sized nuclei (h) in depth bins (mean ± SEM of fields). In KO, nuclei become larger and spermatid-sized nuclei are nearly absent above the basal layer. **(i)**Enlarged 40× views of tubule centres, red (SOX9); green (*γ*H2AX); blue (DAPI). WT: densely packed small spermatid nuclei and SOX9-immunoreactive bodies. KO: large nuclei and no spermatids. Scale bars, 20 µm.

Radial profiles from the BM towards the lumen located the loss (Fig. 4g, h). In WT, median nuclear area stayed at about 22 µm² at every depth, and small nuclei made up about 40% of nuclei throughout. In KO, nuclear size *increased* towards the lumen, from 35 to 46 µm² (Fig. 4g), and small nuclei were nearly absent (< 10%) beyond the basal layer (Fig. 4h). Accordingly, WT tubule centres were packed with small spermatid nuclei and SOX9-immunoreactive bodies, whereas KO tubule centres contained large, spermatocyte-sized or multinucleated cells (Fig. 4i). Germ-cell loss is therefore confined to the adluminal compartment, beyond the stage of early spermatocytes.

### A five-fold expanded interstitium of enlarged, autofluorescent cells

In 20× fields, which were imaged at identical exposure in both genotypes, we plotted the cytoplasmic *γ*H2AX-channel intensity of each interstitial cell against its SOX9-channel intensity (Fig. 5a, b). In WT (723 cells), most cells lay near the origin, whereas in KO (5,736 cells) the two channels were tightly coupled along the identity line. The fraction of cells above the co-brightness threshold in both channels doubled from 25.2% to 50.1% per field (Fig. 5h; +24.5 points, 95% CI 16.3–32.8, *P*_adj_ < 0.001) and did so at every threshold between 30 and 85 (Fig. S3b); because this metric uses 20× fields only, animal-level bounds could not be computed. At 40×, the KO interstitium consisted of continuous sheets of large, rounded to polygonal cells with foamy, finely granular cytoplasm that was bright in both channels (Fig. 5i). Matched emission through two filter sets identifies this signal as intrinsic autofluorescence with the properties of lipofuscin, the lysosomal residue of oxidised protein and lipid [21,22], which in testicular interstitial cells emits from 425 to 700 nm and also occurs at a lower level in normal testes [23].

**Figure 5.**
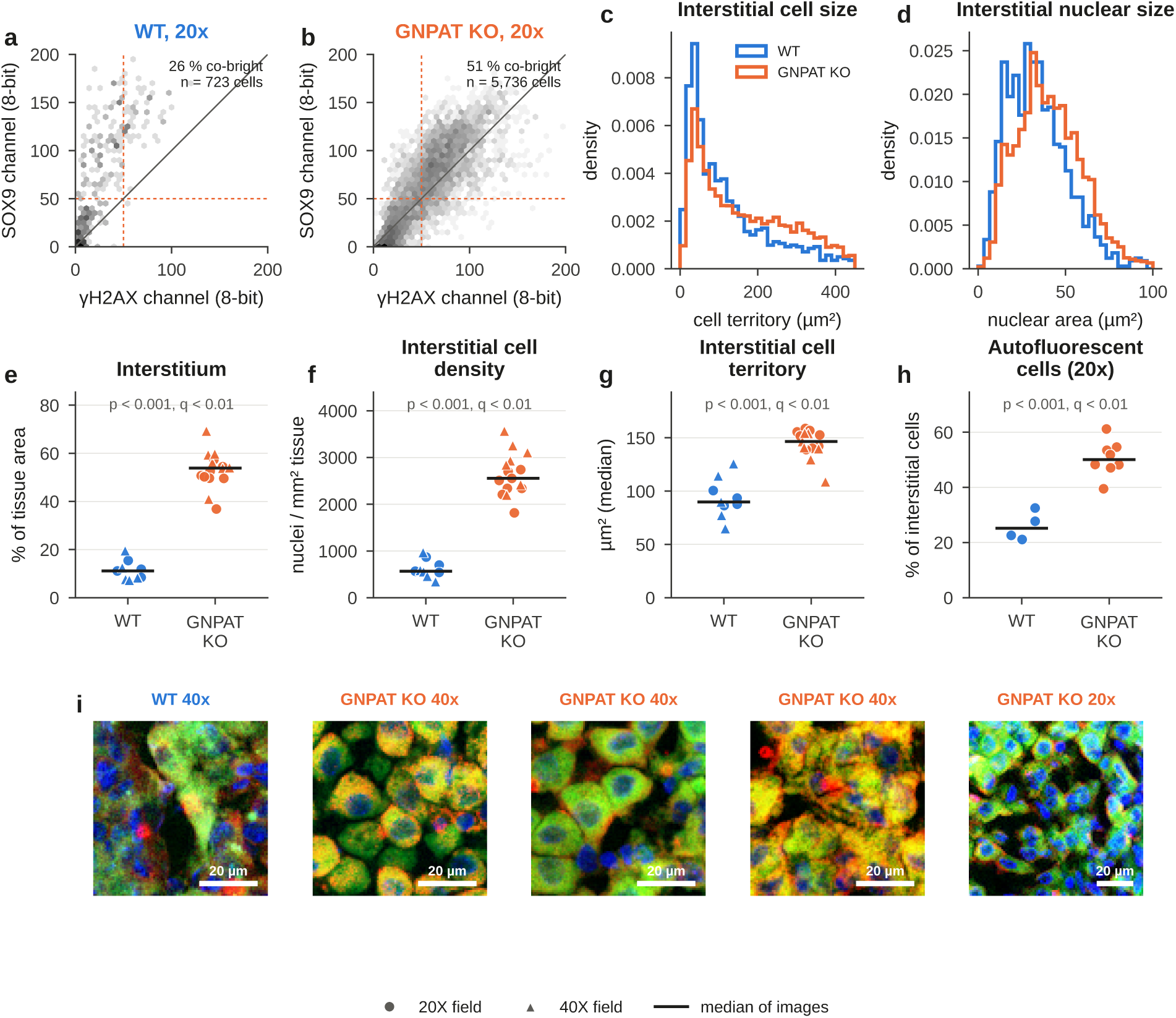
A five-fold expanded interstitium of enlarged, autofluorescent cells. **(a, b)** Mean cytoplasmic intensity in the *γ*H2AX channel (x) vs. the SOX9 channel (y) for every interstitial cell in the 20× fields of WT (a; 723 cells) and KO (b; 5,736 cells), imaged at identical exposure. Grey shading, log-scaled point density; dashed lines, co-brightness threshold (50, 8-bit) in each channel; solid line, identity. The percentage of cells above both thresholds is given in each panel. **(c, d)** Distributions of interstitial cell territory (c) and interstitial nuclear area (d), all cells pooled per genotype. **(e–h)** Field-level interstitium area as % of tissue (e), interstitial nuclei per mm² of tissue (f), median cell territory (g) and proportion of autofluorescent interstitial cells in 20× fields (h). Symbols as in Fig. 3. **(i)** Enlarged views of the interstitium, red (SOX9); green (*γ*H2AX); blue (DAPI): one WT 40× field and four KO fields (three at 40×, one at 20×), showing sheets of large cells with granular, foamy cytoplasm that is bright in both channels. Scale bars, 20 µm. Interstitial cell packing per area of interstitium is shown in Fig. S4e.

The expanded interstitium (53.9% vs. 11.1% of the tissue; Fig. 5e) contained 4.5 times more nuclei per mm² of tissue (2,554 vs. 567; Fig. 5f; *P*_animal_ = 0.003). Interstitial cell territories were 1.6 times larger (median 147 vs. 90 µm²; Fig. 5g; *P*_animal_ = 0.017), with a long tail above 200 µm² (Fig. 5c), and interstitial nuclei were also larger (median 39.7 vs. 32.3 µm²; Fig. 5d). The WT territory matched the mean profile area of a rat Leydig cell (about 91 µm²) [24] (Fig. S4d), which supports the segmentation. Per area of interstitium, however, cell packing did not increase (4,948 vs. 5,633 nuclei per mm²; field-level *P* = 0.04, *P*_animal_ = 0.38; Fig. S4e). The higher density per tissue area therefore reflects a larger compartment filled with larger cells rather than tighter packing.

#### SOX9 and *γ*H2AX immunofluorescence

Both markers gave mainly perinuclear or cytoplasmic signal in tubular nuclei of both genotypes (Fig. S5; Table 1). The median nuclear/perinuclear log_2_ *γ*H2AX enrichment was −0.31 in WT and −0.38 in KO (Fig. S5a; *P*_adj_ = 0.008, *P*_animal_ = 0.22), and *γ*H2AX-enriched nuclei were rare in both genotypes (1.7% vs. 3.7%; Fig. S5b), a difference that reversed sign at a lower cut-off (Fig. S3c). Tubular *γ*H2AX foci did not differ (0.08 vs. 0.04 per nucleus; Fig. S5c; *P*_adj_ = 0.44; Poisson mixed-model rate ratio 0.66, 95% CI 0.29–1.48), interstitial foci were fewer in KO (0.90 vs. 0.35; Fig. S5d; *P*_adj_ = 0.03), and neither foci metric was consistent between magnifications (Fig. S2d). Adluminal KO nuclei were, if anything, less *γ*H2AX-enriched than WT nuclei (Fig. S5h, i). The arrested KO epithelium therefore showed no increase in DNA-damage signalling, although the canonical meiotic *γ*H2AX patterns were not resolved in WT sections (Fig. S5k, left).

SOX9 enrichment of tubular nuclei was −1.09 in WT and −1.41 in KO (Fig. S5e; *P*_adj_ = 0.07), and only 3–4% of tubular nuclei were SOX9-nuclear in either genotype (Fig. S5f, k, right), so individual Sertoli nuclei could not be counted reliably. Nuclear-enriched SOX9 cells within 10 µm of the BM, where Sertoli nuclei sit, were nevertheless 2.4 times more frequent per tubule area in KO (1.9 vs. 0.8 per 10^4^ µm²; Fig. S5g; *P*_adj_ = 0.008, *P*_animal_ = 0.16), independent of the width of the basal band (Fig. S3d), while the radial SOX9 profile was lower throughout the KO epithelium (Fig. S5j). This suggestive result is consistent with relative enrichment of Sertoli cells in a shrunken epithelium.

### Loss of anucleate SOX9-immunoreactive round bodies

The SOX9 channel also showed compact (median equivalent diameter 4.2 µm), saturated, round bodies without DAPI or *γ*H2AX-channel signal, inside WT tubules and in the KO interstitium (Fig. 6g). Because they contain no DNA, they cannot be SOX9-positive nuclei; we refer to them as SOX9-immunoreactive bodies, an operational term that does not imply SOX9 content. They were 2.2-fold rarer in KO, at 125 vs. 292 per mm² of tissue (Fig. 6a; *P*_adj_ < 0.001), 108 vs. 262 per mm² of tubule (Fig. 6b; *P*_adj_ < 0.001) and 126 vs. 346 per mm² of interstitium (Fig. 6c; *P*_adj_ = 0.008), with a negative-binomial rate ratio of 0.45 (95% CI 0.35–0.57; *P*_animal_ = 0.077). Size distributions were almost identical between genotypes (453 WT and 368 KO bodies; Fig. 6d), so the same class of object was counted, but KO bodies lay closer to the BM (median depth 11 vs. 23 µm; Fig. 6e), and only 43% of them, against 90% in WT, were intratubular (Fig. 6f). Their size, position, lack of DNA and dependence on spermatid production fit residual bodies, the cytoplasm shed by elongating spermatids and phagocytosed by Sertoli cells [25,26].

**Figure 6.**
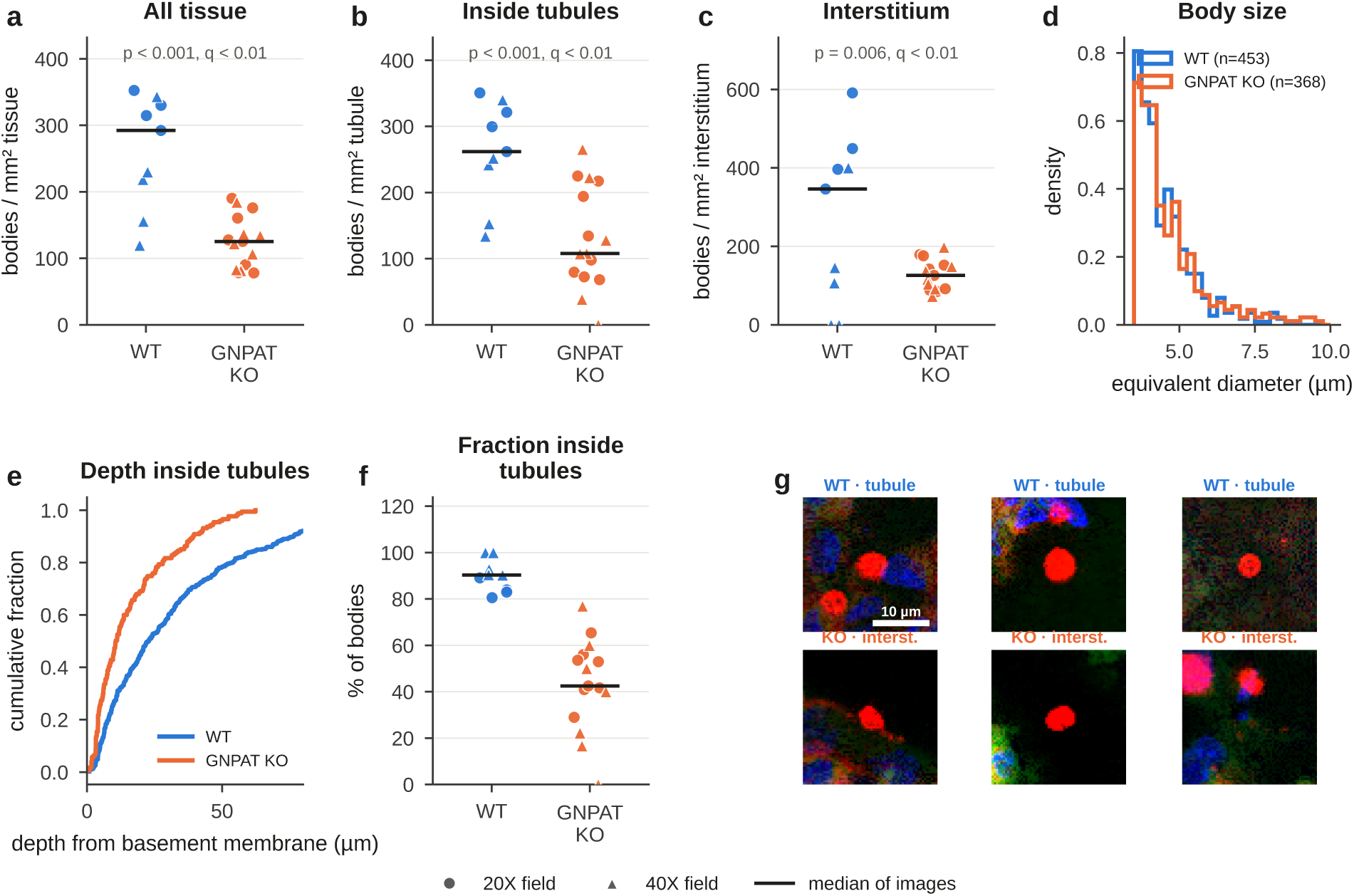
Loss of anucleate SOX9-immunoreactive round bodies. **(a–c)** Density of SOX9-immunoreactive bodies per mm² of tissue (a), per mm² of tubule (b) and per mm² of interstitium (c) in each field. Symbols as in Fig. 3; *p* and *q* (*P*_adj_) printed as in Fig. 3. **(d)** Size distribution (equivalent diameter) of all detected bodies (WT n = 453, KO n = 368). **(e)** Cumulative distribution of the depth of intratubular bodies from the BM. **(f)** Proportion of bodies located inside tubules per field. **(g)** Examples of detected bodies, red (SOX9); green (*γ*H2AX); blue (DAPI): top row, inside WT tubules; bottom row, in the KO interstitium. The bodies are compact, saturated in the SOX9 channel, and negative for DAPI and the *γ*H2AX channel. Scale bar, 10 µm.

### An integrated and robust phenotype

Expressed as standardised KO − WT differences, 18 of the 21 field-level metrics differed at a false-discovery rate below 5% (Fig. 7a). Six also held in the conservative animal-level analysis under the least favourable grouping of fields: tubule area, interstitium area, interstitial nuclear density, interstitial cell territory, tubular nuclear area and the proportion of spermatid-sized nuclei (Table 1). A principal component analysis of the 20 metrics available at both magnifications separated every KO field from every WT field along PC1, which explained 50% of the variance (Fig. 7b) and contrasted interstitial expansion, interstitial cell size and tubular nuclear size with tubule area, SOX9-immunoreactive bodies and small nuclei (Fig. 7c). Effect sizes computed separately in 20× and 40× fields agreed for all architectural, epithelial and interstitial metrics (Fig. S2d), and the loss of spermatid-sized nuclei, the excess of autofluorescent cells and the excess of basal SOX9-nuclear cells held over wide ranges of their analysis thresholds (Fig. S3a, b, d). Only the *γ*H2AX metrics depended on magnification or cut-off (Figs. S2d and S3c). The resulting working model is shown in Fig. 7d.

**Figure 7.**
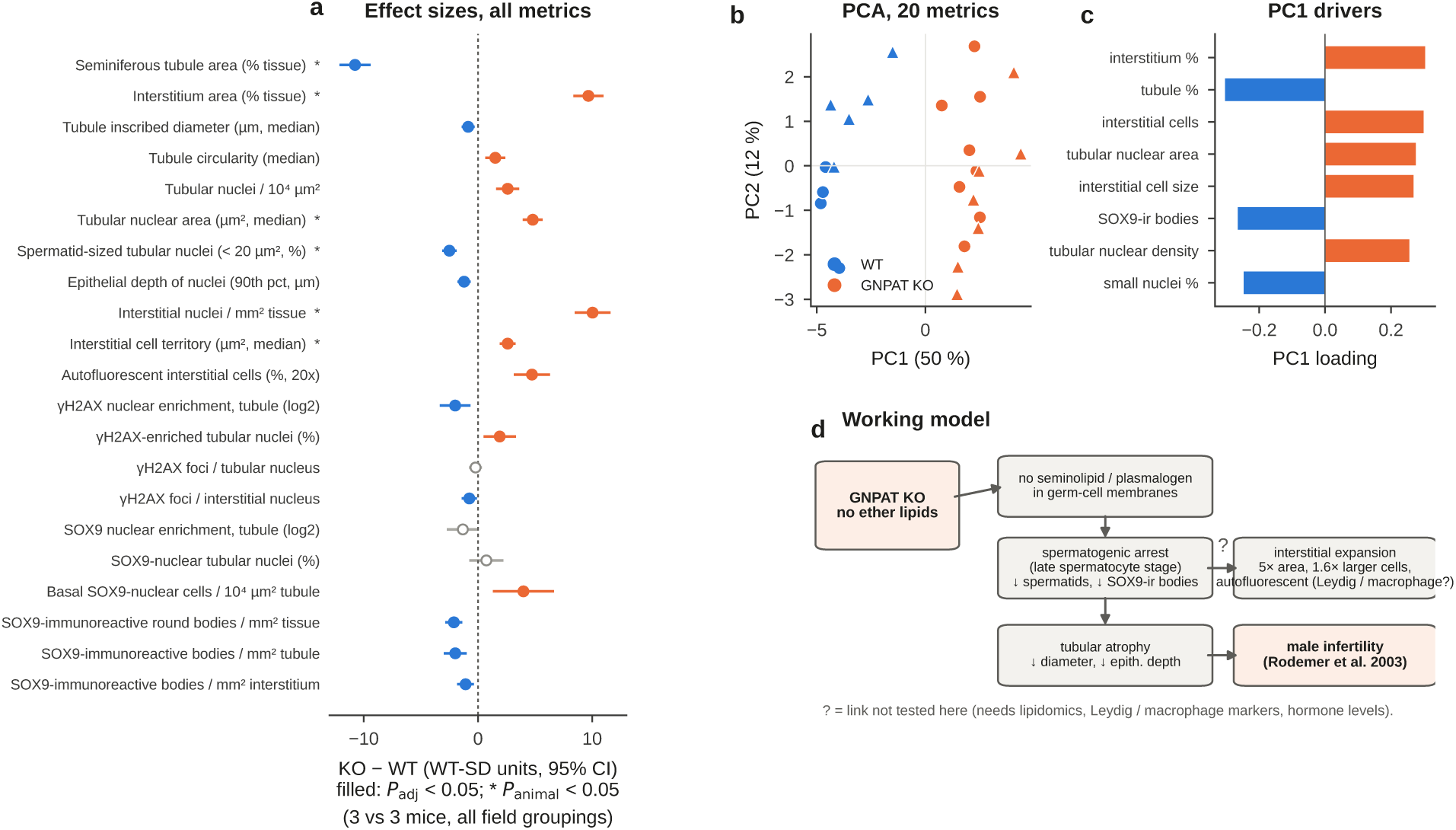
Integrated phenotype and working model. **(a)** Standardised KO − WT difference (in units of the WT standard deviation, 95% CI) for all 21 field-level metrics. Orange, higher in KO; blue, lower in KO. Filled symbols, *P*_adj_ < 0.05; open symbols, *P*_adj_ ≥ 0.05; asterisk, significant in the conservative animal-level analysis (*P*_animal_ < 0.05, i.e. *P* < 0.05 for every grouping of fields into 3 + 3 mice). **(b)** Principal component analysis of the 20 metrics available at both magnifications (*z*-scored). Each symbol is one field (circles, 20×; triangles, 40×); WT and KO separate completely along PC1 (50% of variance). **(c)** The eight metrics with the largest PC1 loadings. **(d)** Working model. GNPAT loss removes seminolipid and plasmalogens from germ-cell membranes, leading to spermatogenic arrest at the late spermatocyte stage. Spermatids and SOX9-ir bodies are lost, and the tubules atrophy. The interstitium expands (five-fold area, 1.6-fold larger, autofluorescent cells of Leydig or macrophage character). “?” marks links not tested here, which need lipidomics, Leydig and macrophage markers, and hormone measurements. Male infertility is from [12].

## Discussion

Two decades after male infertility was reported in ether-lipid-deficient mice [12], this study gives the first quantitative, fluorescence-based description of the *Gnpat*-null testis. It confirms the known germ-cell phenotype with effect sizes and animal-level statistics and adds three observations: a five-fold expansion of the interstitium with enlarged cells, a doubling of autofluorescent interstitial cells, and loss of anucleate SOX9-immunoreactive bodies (Fig. 7d; Table S1).

Our measurements are of the size expected from published models (Fig. S4). The reduction in tubule diameter of about 30% lies with the 40% reported on semithin sections of an independently generated *Gnpat*-knockout line [12] and with germ-cell *Pex13* knockouts (−30%) [17] and a Sertoli-driven meiotic block (−39%) [27] (Fig. S4a). The 74% loss of spermatid-sized nuclei is less extreme than transcript-based measures in *Far1*- and germ-cell *Pex13*-null testes [10,17], as expected for a size-based classification, and the WT interstitial fraction approaches the 16% measured stereologically in normal rat testis [24] (Fig. S4c). A phenotype matrix built from 265 quote-verified evidence items in 59 studies (Fig. S6; Table S2) shows that tubular atrophy, meiotic-to-spermatid arrest and multinucleated cells are shared by every model that removes ether lipids (*Gnpat*, *Agps*, *Far1*) or seminolipid (*Cgt*, *Cst*) and by germ-cell peroxisome deficiency (*Pex13*, *Pex5*) [8,10,12,15,17,18]. Interstitial lipid or pigment, by contrast, had been reported only qualitatively and never as autofluorescence [17,28]. The literature strongly supports the atrophy, the arrest, the loss of residual bodies and the pigment, whereas the interstitial expansion, the SOX9 result and the lipid mechanism draw challenges (Fig. S7), which we address below; Fig. S8 separates established, measured and hypothetical steps of the resulting model.

### Which ether lipid drives the germ-cell arrest?

*Gnpat* deletion removes plasmalogens and seminolipid together, but several lines of evidence point to seminolipid as the main germ-cell lesion. Its synthesis needs GNPAT, AGPS and FAR1, and loss of any of these enzymes, or of CGT or CST downstream, arrests spermatogenesis in the same pachytene-to-metaphase I window with multinucleated cells [8,10,11,15]. The seminolipid defect is germ-cell-autonomous [8], and its proposed mechanism, failed cytokinesis at intercellular bridges [10], fits the multinucleated cells seen here and before [12]. Conversely, *Peds1*-null mice, which lack plasmalogens but retain alkyl ether lipids, are fertile [6], although testis histology and seminolipid were not examined in that preprint. Plasmalogen loss may still contribute: ether-lipid deficiency disrupts blood–testis-barrier remodelling [14], and our radial profiles place the germ-cell loss exactly behind that barrier.

### The arrest is not a DNA-damage arrest

Meiotic arrests caused by unrepaired double-strand breaks or failed synapsis raise autosomal *γ*H2AX several-fold, by 5- to 7-fold in *Trip13* mutants and germ-cell *Pex5* knockouts [18,29,30] (Fig. S4b). We saw no such increase, which fits a lipid-dependent block during the meiotic divisions or cytokinesis, as in *Cst*-null mice [8], rather than a recombination defect. Two limitations temper this conclusion: section-level counts can fall when meiotic cells are lost even if per-cell repair is impaired [18], and our antibody did not resolve the canonical meiotic patterns [31]. Spermatocyte spreads stained for SYCP3 and *γ*H2AX remain the decisive test [32].

### Sertoli cells and residual bodies

The higher density of basal SOX9^+^ cells is best read as relative enrichment, because adult Sertoli cells are post-mitotic [33], their number does not change when germ cells are lost [34,35], and it is normal in germ-cell *Pex5* knockouts [18]. The loss of residual bodies follows from the arrest: Sertoli cells phagocytose residual bodies at spermiation and digest them in lysosomes [25,26], so fewer spermatids mean fewer bodies. Sertoli cells also take up seminolipid from phagocytosed germ cells, and impaired seminolipid degradation causes lysosomal storage [36], a link between germ-cell lipid and phagocyte lysosomes that may bear on the interstitial pigment.

### A new interstitial phenotype: expansion, hypertrophy and pigment

The most striking new finding is the extent of the interstitium. Its five-fold expansion reflects a larger compartment of larger cells rather than tighter packing. In adult mice, depleting germ cells alone does not change Leydig-cell number [34,35], whereas in human non-obstructive azoospermia Leydig cells are hypertrophic and lipid-laden and macrophages increase [28]. True Leydig hyperplasia occurs when Sertoli function or endocrine signalling is disturbed [37,38], and it has been described in *Gnpat*-knockout mice [12] and after germ-cell *Pex13* deletion [17]. Our data are therefore consistent with Leydig hypertrophy, with or without some hyperplasia; a stereological count per testis or a per-testis *Hsd3b1* transcript standard is needed to decide [35,39]. Resident macrophages, which are closely associated with Leydig cells and easily mistaken for them, could also contribute [40,41].

Two non-exclusive sources of the pigment are plausible. The first is phagocytic: macrophages that clear apoptotic germ cells accumulate lysosomal residue, and in azoospermic men they carry many more lipofuscin bodies than in controls [22,28]. The second is a cell-intrinsic defect in lipid handling. Plasmalogen-deficient cells transport cholesterol poorly to the endoplasmic reticulum [19,20], steroidogenic Leydig cells handle large cholesterol fluxes through autophagy [42,43], peroxisome-deficient Leydig cells store large lipid droplets [17], and testicular lipid inclusions with testicular insufficiency are common in X-linked adrenoleukodystrophy [44]. Because ether-lipid deficiency reduces rather than increases lipid droplets in other tissues [16], a trafficking or lysosomal defect that produces lipofuscin is more likely than simple lipid overload. Sudan Black B, filipin and BODIPY staining with Leydig and macrophage markers can distinguish these possibilities [23].

### Implications for fertility and therapy

The interstitial phenotype raises the possibility that ether-lipid deficiency affects male fertility through both a germ-cell route and an interstitial, possibly endocrine, route. The gonadal phenotype of RCDP in humans is undocumented [7], and *GNPAT* is not among the genes confidently linked to human male infertility [45]. Men with milder RCDP or secondary plasmalogen deficiency might nevertheless have impaired testicular function, as do most men with adrenomyeloneuropathy, another peroxisomal disorder [44]. Alkylglycerols enter the pathway after the peroxisomal steps, restore plasmalogens in peripheral organs and, in *Pex7*-null mice, restored testicular plasmalogens and spermatocytes [4,16]. Because they also supply the alkyl backbone of seminolipid, a rescue experiment could ask whether the interstitial pigment regresses along with the germ-cell arrest.

### Conclusion

Ether-lipid deficiency does more than arrest spermatogenesis. It remodels the whole testis, shrinking the seminiferous epithelium and expanding an interstitium of enlarged, pigment-laden cells. Together with the published models, our data favour seminolipid loss as the germ-cell lesion and identify the interstitium as a second, previously unmeasured compartment of ether-lipid-related male infertility.

## Materials and methods

### Animals

Heterozygous *Gnpat* knockout mice (*Gnpat*^+/−^) on a C57BL/6J background were generated and supplied by Cyagen Biosciences (contract no. GEM-KOAIP-230630-CAL-03; *Gnpat* knockout line EK5264, founder line 2-1766). Two male and two female *Gnpat*^+/−^ breeders (born 16 August 2023) were received on 16 October 2023 and intercrossed to obtain wild-type (WT; *Gnpat*^+/+^) and knockout (KO; *Gnpat*^−/−^) littermates. Male WT and KO littermate mice (n = 3 per genotype), all on the C57BL/6J background, were analysed at 12 weeks of age. [Genotyping method; housing conditions (temperature, light cycle, food and water *ad libitum*).] All animal experiments were approved by the Ethics Committee for Animal Experiments of the Institute of Rheological Functions of Food [approval number] and were carried out in accordance with the institutional guidelines. The study is reported in line with the ARRIVE guidelines.

### Tissue collection, fixation and cryosectioning

Mice were deeply anaesthetised with [anaesthetic, dose, route] and euthanised by [method]. Both testes were dissected, weighed [if recorded] and immersion-fixed in freshly prepared 4% paraformaldehyde (PFA) in 0.1 M phosphate-buffered saline (PBS, pH 7.4) for [24 h] at 4 °C. Fixed testes were rinsed in PBS and cryoprotected at 4 °C in 15% and then 30% (w/v) sucrose in PBS, each until the tissue sank. They were embedded in optimal cutting temperature (OCT) compound (Tissue-Tek, Sakura Finetek), frozen on dry ice [or in isopentane cooled on dry ice] and stored at −80 °C. Cryosections were cut at 15 µm on a cryostat [model] at −20 °C, mounted on positively charged glass slides (e.g. Superfrost Plus), air-dried for 30 min at room temperature (RT) and stored at −80 °C until staining.

### Immunohistochemistry

Sections from WT and KO testes were stained in parallel in the same session with identical reagents. Slides were brought to RT, air-dried for 10 min and washed three times for 5 min in PBS to remove OCT. [If performed: heat-induced antigen retrieval was carried out in 10 mM sodium citrate, pH 6.0, at 95 °C for 10 min, followed by cooling to RT.] Sections were permeabilised with 0.3% Triton X-100 in PBS for 15 min and blocked for 1 h at RT with 5% normal goat serum and 1% bovine serum albumin (BSA) in PBS containing 0.1% Triton X-100. [If used: endogenous mouse IgG was blocked with a mouse-on-mouse blocking reagent before application of the mouse primary antibody.] Sections were then incubated overnight at 4 °C in a humidified chamber with a SOX9 recombinant rabbit monoclonal antibody (clone 7H13L8; Invitrogen, Thermo Fisher Scientific; [cat. no. 702016]; [1:200]) and a mouse monoclonal phospho-histone H2A.X (Ser139) (*γ*H2AX) antibody (Cell Signaling Technology; [clone 2F3, cat. no. 80312]; [1:400]), both diluted in blocking buffer. After three 10-min washes in PBS containing 0.1% Tween-20 (PBST), sections were incubated for 1 h at RT in the dark with goat anti-rabbit IgG (H+L) conjugated to Alexa Fluor 594 ([Invitrogen, A-11012]; 1:500) to detect SOX9 and goat anti-mouse IgG (H+L) conjugated to Alexa Fluor 488 ([Invitrogen, A-11001]; 1:500) to detect *γ*H2AX. After three further PBST washes, nuclei were counter-stained with 4’,6-diamidino-2-phenylindole (DAPI; 1 µg/mL in PBS) for 10 min, and sections were rinsed in PBS and mounted under coverslips in antifade medium ([e.g. ProLong Gold, Invitrogen]). Slides were cured overnight at RT in the dark and stored at 4 °C until imaging. [Secondary-antibody-only control sections, with the primary antibodies omitted, were processed in parallel — state whether performed.] Because the *γ*H2AX primary antibody was raised in mouse, the anti-mouse secondary antibody can also bind endogenous mouse immunoglobulin, which is abundant in the interstitium and in vessels. This does not affect our autofluorescence measure, which requires matched signal in both the *γ*H2AX and the SOX9 channel, the latter being detected with an anti-rabbit reagent.

### Image acquisition

Images were acquired on a Keyence BZ-X fluorescence microscope [model, e.g. BZ-X810] with PlanApo 20×/0.75 and 40×/0.95 objectives [confirm manufacturer], giving pixel sizes of 0.8 and 0.4 µm, respectively. DAPI was imaged with a DAPI filter cube (excitation 360/40 nm, emission 460/50 nm), the Alexa Fluor 488 (*γ*H2AX) signal with a GFP cube (470/40 nm, 525/50 nm) and the Alexa Fluor 594 (SOX9) signal with a TRITC cube (545/25 nm, 605/70 nm) [confirm cube specifications]; each channel is named after the antibody imaged in it. For each field, a z-stack of 22–46 planes at 0.2–0.5 µm steps was fused into one extended-depth-of-field image per channel with the BZ-X Analyzer software and exported as a three-channel 8-bit

OME-TIFF of 916 × 687 pixels (733 × 550 µm at 20×; 366 × 275 µm at 40×). Exposure times were read from the instrument metadata (Fig. S2a, b). Because the *γ*H2AX-channel exposure differed between genotypes at 40×, all marker readouts are within-image ratios, and the autofluorescence metric uses 20× fields only, where exposures were identical. Fields were selected to contain tubules and interstitium and to avoid folds and edge artefacts; raw files were opened read-only and verified against MD5 checksums.

### Image analysis

All image analysis was automated in Python, with every length and area defined in micrometres and converted with each image’s calibration; full parameters are given in the Supplementary methods. Tubule, interstitium and background were assigned by a random-forest pixel classifier (scikit-learn [46]) trained on sparse annotations in eight fields of both genotypes and objectives, using 102 multiscale features of the three channels, a *γ*H2AX/SOX9-channel log-ratio and a co-brightness map. Touching tubule profiles were separated by a distance-transform watershed, and each profile’s area, inscribed diameter and circularity were recorded. Nuclei were segmented in the DAPI channel with the Cellpose *nuclei* model [47] (diameter 7.5 µm), assigned to a compartment and, in tubules, given a depth from the BM; tubular nuclei were classed as small (< 20 µm², spermatid-sized), intermediate or large (> 40 µm²). Marker enrichment was calculated as *E* = log_2_[(*Ī* _nucleus_ + 1)/(*Ī* _ring_ + 1)] with a 2-µm perinuclear ring, so that it is independent of exposure; *γ*H2AX foci were detected with a difference-of-Gaussians filter. Interstitial cell territories were grown from each interstitial nucleus by watershed, and a cell was scored as autofluorescent when its mean cytoplasmic intensity was at least 50 (8-bit) in both the *γ*H2AX and the SOX9 channel. SOX9-immunoreactive bodies were detected as compact, round, isolated objects specific to the SOX9 channel. Twenty-one metrics were derived per field, which served as the unit of analysis.

### Statistical analysis

For each field-level metric we fitted a linear model with genotype and magnification as predictors and report the KO − WT difference with its 95% CI; *P* values were adjusted for the 21 metrics with the Benjamini– Hochberg procedure [48] (*P*_adj_). Tubule-level diameter and nucleus-level nuclear area and marker enrichment were analysed with linear mixed models with a random intercept per field (lme4 [49]; lmerTest [50]), *γ*H2AX foci with a Poisson mixed model with an observation-level random effect, and counts of SOX9-immunoreactive bodies with a negative-binomial model with an offset for tissue area (MASS). Because the fields came from three mice per genotype and treating them as independent would be pseudoreplication [51], we also enumerated every balanced grouping of fields into animals (35.3 million combinations), averaged fields within each animal and performed a two-sample *t*-test (3 vs. 3 mice; df = 4) for each grouping. The largest *P* value over all groupings is reported as the conservative animal-level *P* value (*P*_animal_), so that *P*_animal_ < 0.05 holds for any assignment of fields to animals. Standardised effect sizes, Hedges’ *g*, consistency between magnifications, threshold sensitivity and a principal component analysis of the *z*-scored metrics are described in the Supplementary methods. All tests were two-sided.

### Literature synthesis

Fifty-eight open-access studies and reviews identified through Europe PMC, together with the original description of the *Gnpat* knockout [12], were analysed for statements relevant to our findings. Each statement was recorded as a verbatim quote, checked programmatically against the full text and classified as supporting, challenging or giving context (Table S2; Supplementary methods).

### Software

Image analysis used Python 3.12 with tifffile 2026.4, NumPy 2.0, SciPy 1.18, scikit-image 0.26, scikit-learn 1.9 [46], Cellpose 3.1 [47], pandas 3.0 and matplotlib 3.11. Statistical analysis used R 4.3.3 with lme4 2.0 [49], lmerTest 3.2 [50] and MASS 7.3.

## Declarations

### Data and code availability

All code and raw or processed data are available from the corresponding authors upon reasonable request.

### Author contributions

N.S. designed the experiments, performed all analyses to generate the data, and developed the computational and mathematical analysis software used to examine the tissue histology quantitatively. A.F. developed the image-analysis software. M.H. designed the experiments. S.M. designed the experiments and helped perform the research. T.F. designed the experiments and helped write the manuscript.

## Supplementary figures

**Figure S1.**
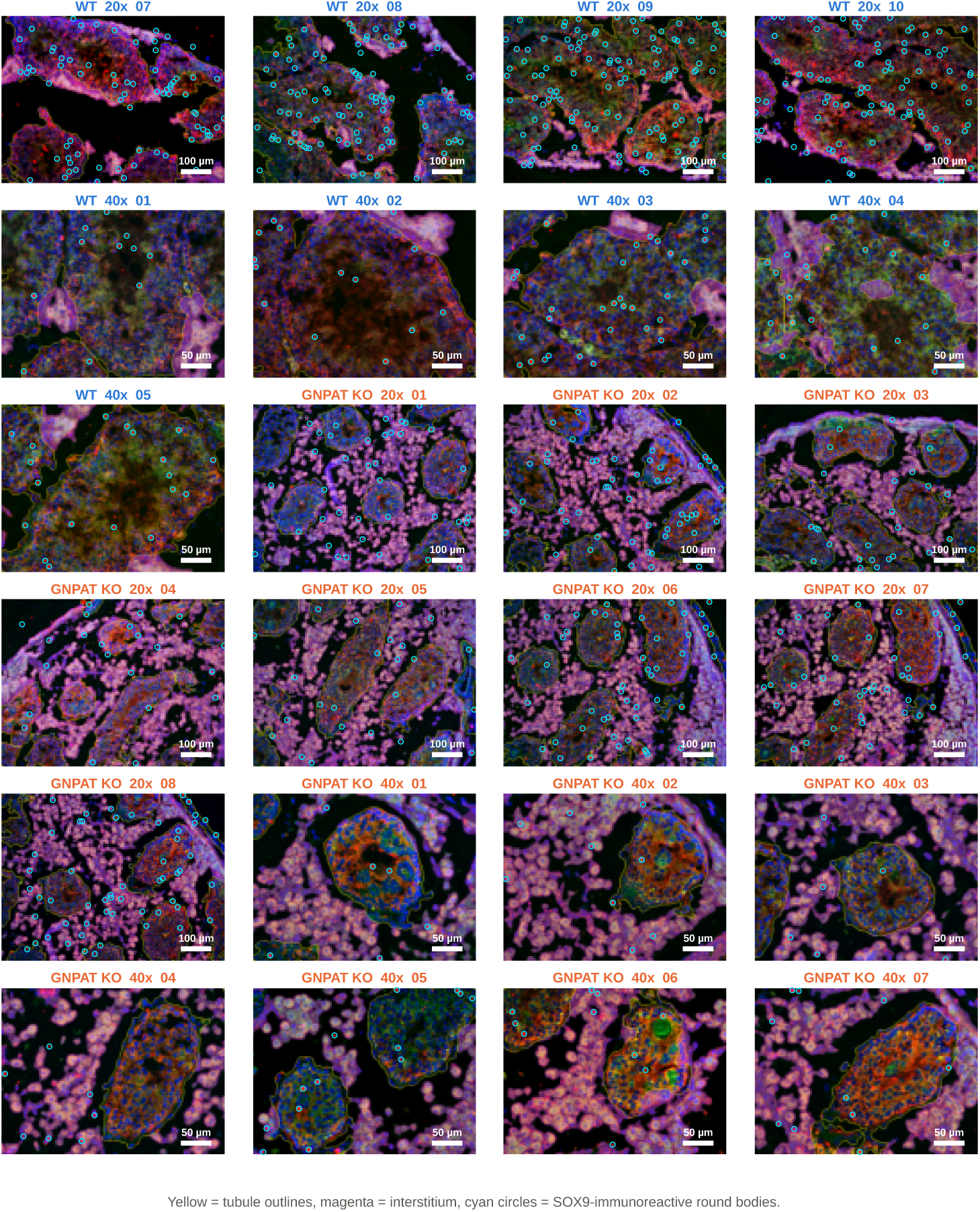
Segmentation quality control for all 24 fields. Each panel is one field, labelled by genotype, magnification and field number. Images show red (SOX9); green (*γ*H2AX); blue (DAPI), with segmentation overlays: yellow lines, tubule outlines; magenta tint, interstitium; cyan circles, detected SOX9-immunoreactive round bodies. Scale bars: 100 µm (20×), 50 µm (40×).

## Supplementary tables

**Table S1.**
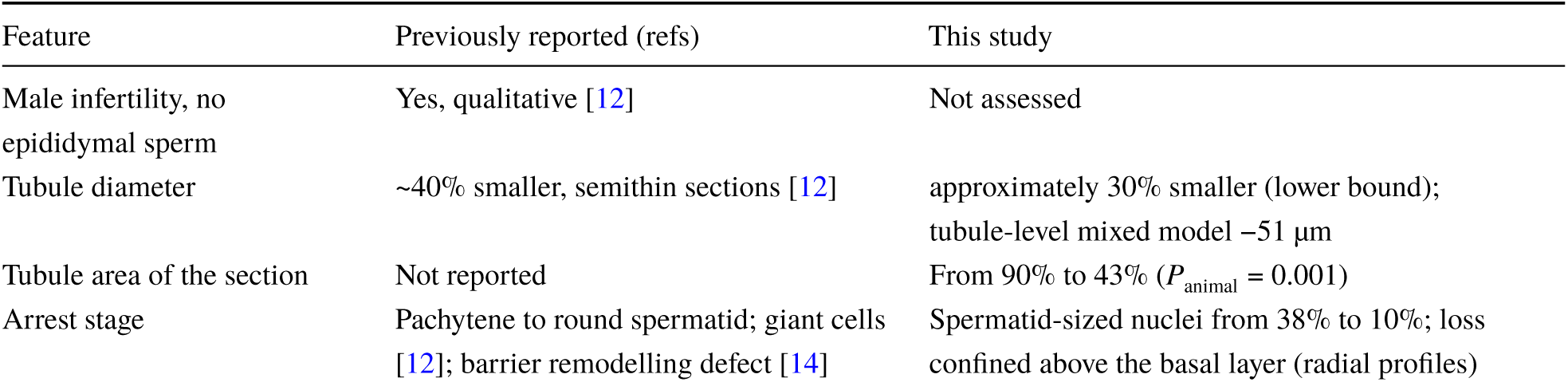
What this study adds to the published description of the *Gnpat*-null testis.

**Table S2.**
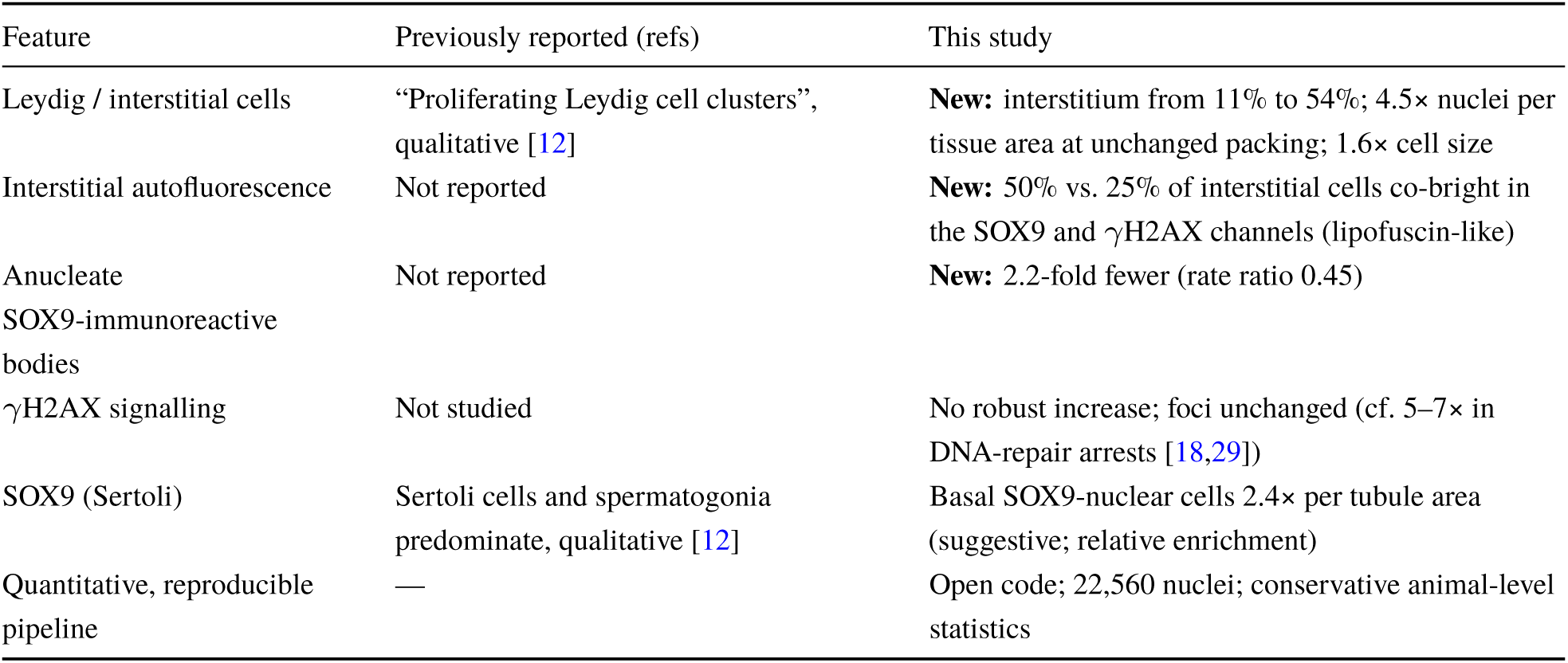
Literature evidence (separate file: Supplementary_Table_S1_literature_evidence.xlsx). The table lists one row per evidence item (265 items from 59 papers), giving the paper, journal, DOI and model; the finding addressed and its relation to our data (supports, challenges or context); a paraphrase and the verbatim quote with its location in the source; and any quantitative values (control, mutant, fold change and unit).

## Supplementary methods

### Image analysis: full parameters

All image analysis was automated in Python. Every length and area was defined in micrometres and converted to pixels with each image’s calibration, so that 20× and 40× fields were processed with identical physical parameters. For classification and display, each channel was rescaled per image between its 1st and 99.5th percentiles within the tissue, which removes differences in exposure and staining intensity between fields. Background (empty slide and tubular lumen) was defined as pixels whose lightly smoothed (σ = 1.5 µm) summed intensity fell below a low, image-adaptive threshold.

Tissue compartments were identified with a random-forest pixel classifier. Each pixel was described by 102 features computed from six input maps: the three normalised channels; the *γ*H2AX/SOX9-channel log-ratio, which is close to zero in autofluorescent cells; the pixel-wise minimum of the *γ*H2AX and SOX9 channels, a co-brightness map; and the tissue mask. For each map we computed the Gaussian-smoothed intensity at five scales (σ = 0.8, 1.6, 3.2, 6.4 and 12.8 µm) and, for σ ≥ 1.6 µm, the gradient magnitude, the scale-normalised Laplacian of Gaussian and the local standard deviation. Training labels were drawn as sparse circular scribbles marking tubule and interstitium in eight fields covering both genotypes and both objectives, with background taken from the background mask. Up to 6,000 pixels per class and image were sampled to train a random-forest classifier (scikit-learn [46]; 200 trees, minimum of 5 samples per leaf, balanced class weights), which returned for every pixel the probabilities of tubule, interstitium and background; these were smoothed with σ = 2 µm.

A pixel was assigned to the tubule compartment when its tubule probability exceeded those of both other classes. The tubule mask was cleaned by morphological opening with a 4-µm disk, objects smaller than 2,500 µm² were removed, and holes were filled except for interstitium-like holes larger than 400 µm². Touching tubule cross-sections were separated by a watershed on the Euclidean distance transform, seeded by distance maxima that lay at least 15 µm from the boundary, at least 70 µm apart and at least 12 µm above the surrounding saddle (h-maxima). Tissue that was neither tubule nor background was classed as interstitium, after removal of objects smaller than 60 µm². For each tubule profile we recorded its area, inscribed diameter (twice the maximum of the internal distance transform), circularity (4π·area/perimeter²) and solidity, and whether it touched the field edge. Profiles with an inscribed diameter below 40 µm or a solidity below 0.7, mostly strips of tunica or oblique cuts, were excluded from tubule-level statistics.

**Figure S2.**
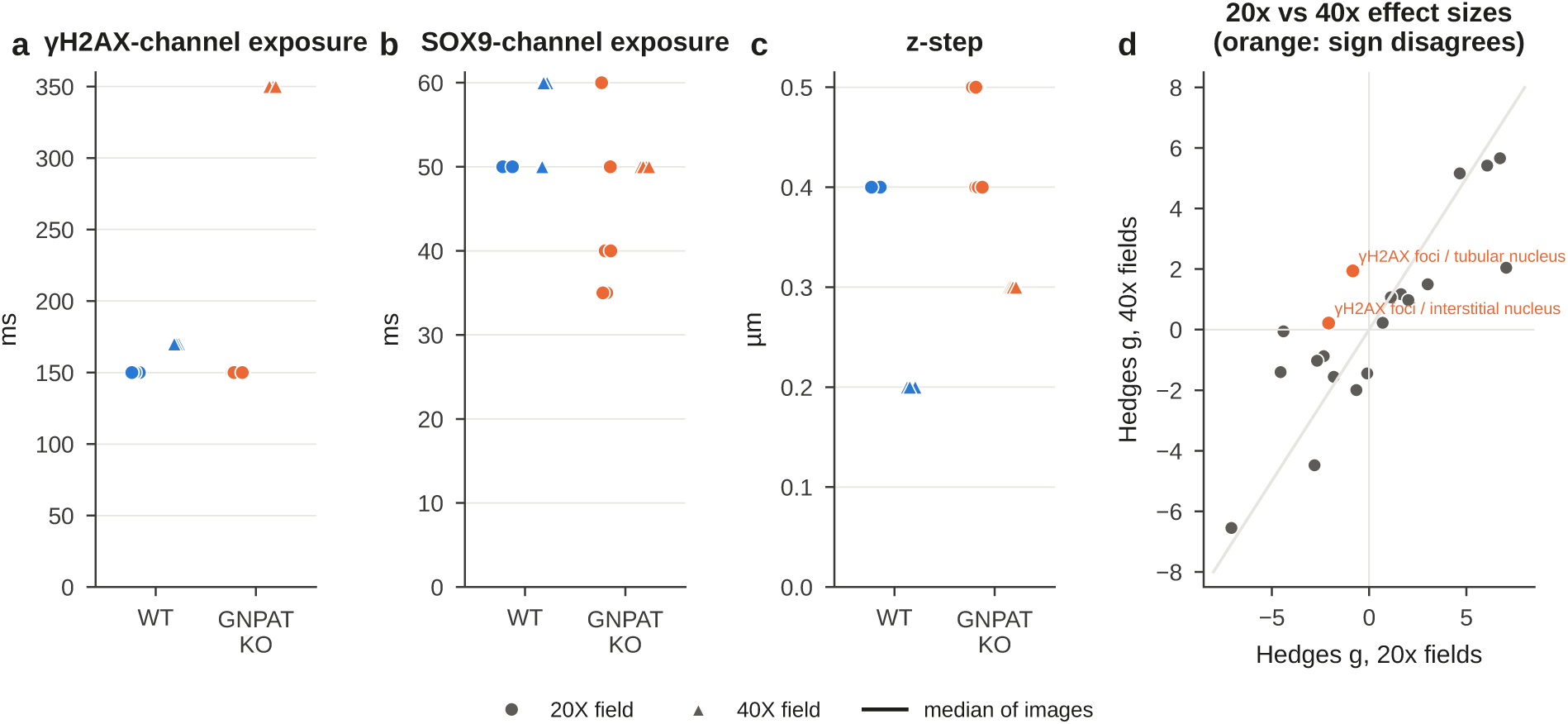
Acquisition parameters and consistency between magnifications. **(a–c)** Exposure time of the *γ*H2AX channel (a) and the SOX9 channel (b), and z-step of the extended-focus stack (c), for every field (circles, 20×; triangles, 40×; bars, medians). *γ*H2AX-channel exposure was identical between genotypes at 20× (150 ms) but differed at 40× (WT 170 ms, KO 350 ms). Raw intensities were therefore not compared between genotypes, and the autofluorescence metric uses 20× fields only. **(d)** Hedges’ *g* (KO vs. WT) computed separately in 20× and 40× fields for each metric. Grey line, identity. Orange symbols mark the only two metrics whose sign differs between magnifications (*γ*H2AX foci per tubular and per interstitial nucleus).

**Figure S3.**
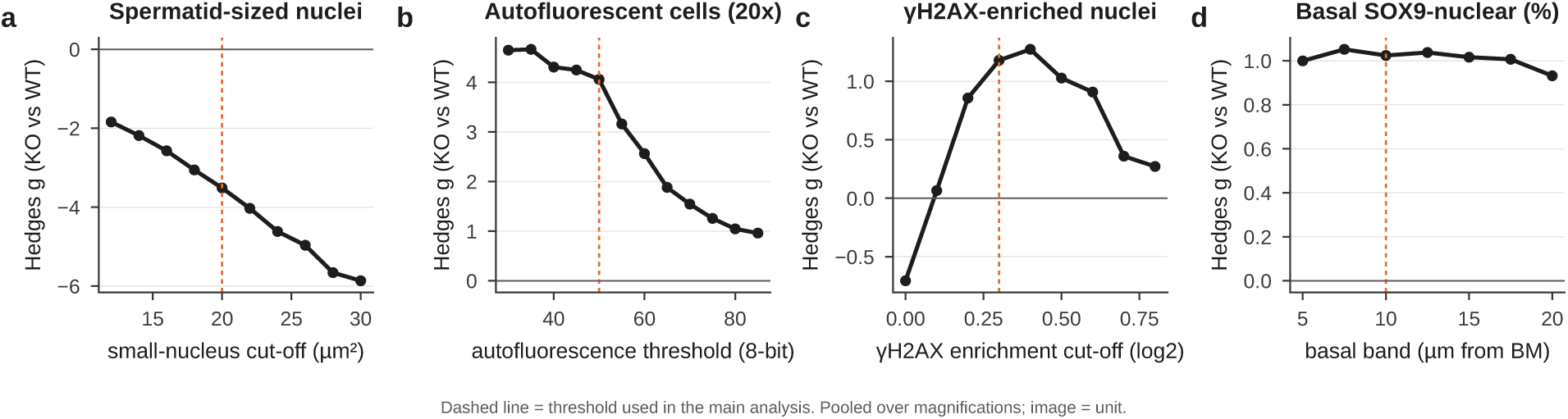
Sensitivity of the main conclusions to analysis thresholds. KO vs. WT Hedges’ *g* (fields pooled over magnifications) recomputed as each threshold is varied. **(a)** Size cut-off defining spermatid-sized tubular nuclei (12–30 µm²). **(b)** Co-brightness threshold for autofluorescent interstitial cells (30–85, 8-bit; 20× fields). **(c)** Enrichment cut-off defining *γ*H2AX-enriched nuclei (log_2_ 0–0.8). The effect changes sign at a cut-off of 0. **(d)** Width of the basal band used to count SOX9-nuclear cells (5–20 µm from the BM). Red dashed lines, values used in the main analysis.

Nuclei were segmented in the DAPI channel with the pretrained Cellpose *nuclei* model [47], a deep-learning instance-segmentation network, using an expected nuclear diameter of 7.5 µm and a flow threshold of 0.5 after scaling DAPI to its 99.8th percentile. Each nucleus was assigned to the compartment containing its centroid. We recorded nuclear area and shape and, for tubular nuclei, the depth from the basement membrane (BM), defined as the Euclidean distance from the nuclear centroid to the tubule boundary. Tubular nuclei were classed as small (< 20 µm², spermatid-sized), intermediate (20–40 µm²) or large (> 40 µm²). Marker enrichment was calculated for each nucleus as *E* = log_2_[(*Ī* _nucleus_ + 1)/(*Ī* _ring_ + 1)], where *Ī* is the mean raw intensity and the ring is a 2-µm perinuclear band that excludes other nuclei and background. *E* > 0 indicates nuclear accumulation and *E* < 0 mainly cytoplasmic signal; because *E* is a ratio within one image, it is independent of exposure. *γ*H2AX foci were detected as local maxima, at least 0.8 µm apart, of a difference-of-Gaussians filter of the *γ*H2AX channel (σ = 0.4 and 1.2 µm) whose robust *z* score (based on the median and median absolute deviation over the image) was at least 4, and were assigned to the nucleus containing them.

**Figure S4.**
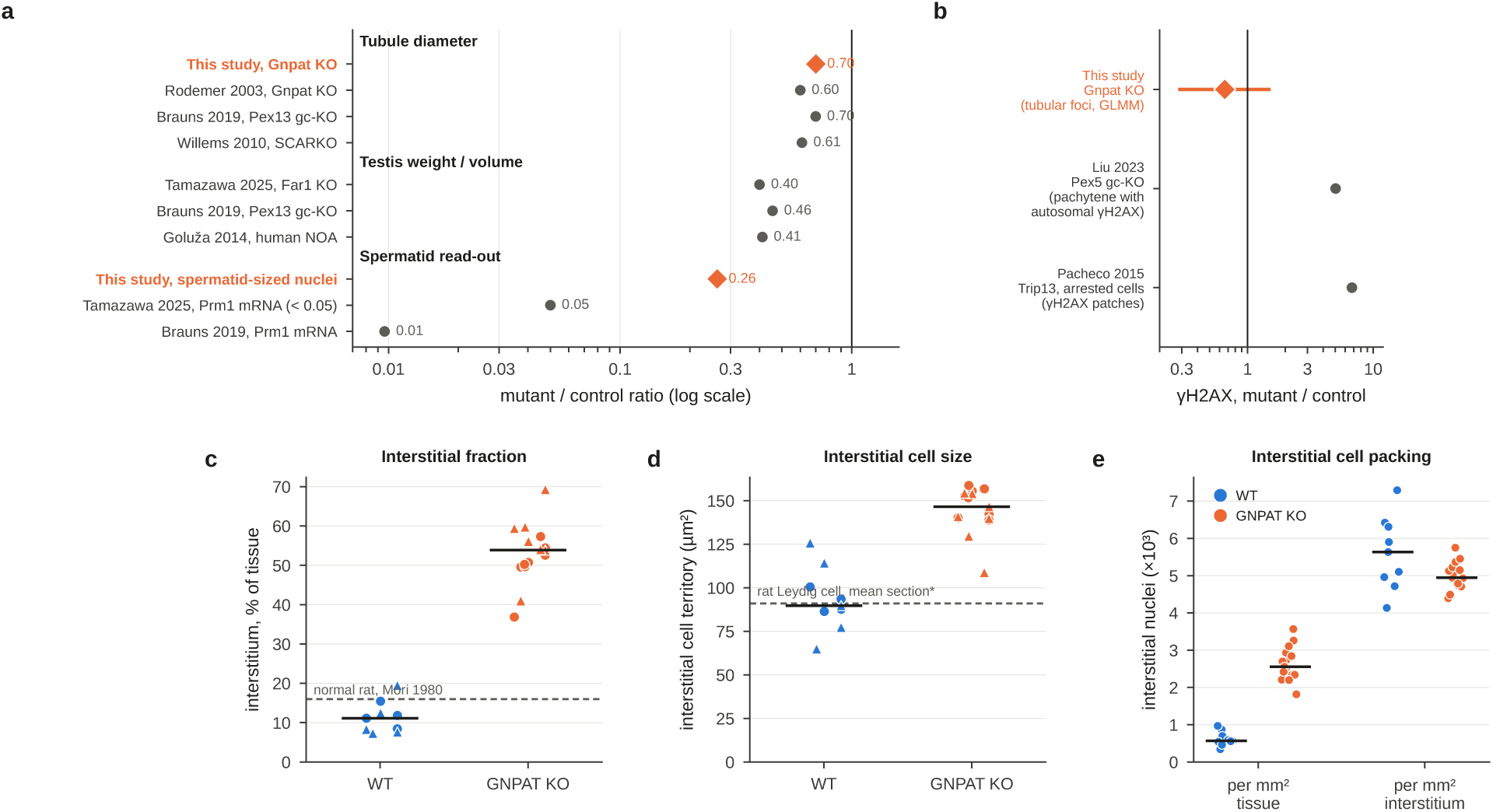
Quantitative benchmarking against published values. **(a)** Mutant/control ratios (log scale) for tubule diameter, testis weight or volume, and spermatid read-outs. Orange diamonds, this study (tubule inscribed diameter; proportion of spermatid-sized nuclei); grey circles, published values, with sources on the left. Our tubule ratio is a lower bound because WT tubules extended beyond the field. **(b)** *γ*H2AX increase in meiotic arrests caused by defective DNA repair (grey; arrested vs. progressing *Trip13*-mutant spermatocytes, and germ-cell *Pex5*knockouts) vs. the tubular *γ*H2AX foci rate ratio in this study (orange; 95% CI from the Poisson mixed model). **(c)** Interstitial area fraction per field; dashed line, stereological value for normal adult rat testis. **(d)** Interstitial cell territory per field; dashed line, mean profile area of a rat Leydig cell, which we derived as (2/3)πr² from the published mean volume of 1,209 µm³. **(e)** Interstitial nuclei per mm² of tissue (left) and per mm² of interstitium (right). The 4.5-fold increase per tissue area disappears when expressed per interstitial area. In (c–e), circles, 20×; triangles, 40×; bars, medians.

Each interstitial nucleus seeded a cell territory, grown by a watershed on the smoothed (σ = 0.8 µm) sum of the *γ*H2AX and SOX9 channels and restricted to the interstitium and to within 8 µm of a nucleus; the cytoplasm was defined as the territory minus the nucleus. A cell was scored as autofluorescent when its mean raw cytoplasmic intensity was at least 50 (8-bit) in both the *γ*H2AX and the SOX9 channel. This measure was computed in 20× fields only, where exposures were identical in both genotypes.

SOX9-immunoreactive round bodies were detected as compact objects specific to the SOX9 channel. Candidate pixels had a normalised SOX9-channel signal exceeding the larger of the normalised *γ*H2AX and DAPI signals by more than 0.35 (after smoothing at σ = 0.4 µm) and a raw SOX9-channel intensity of at least 180.

**Figure S5.**
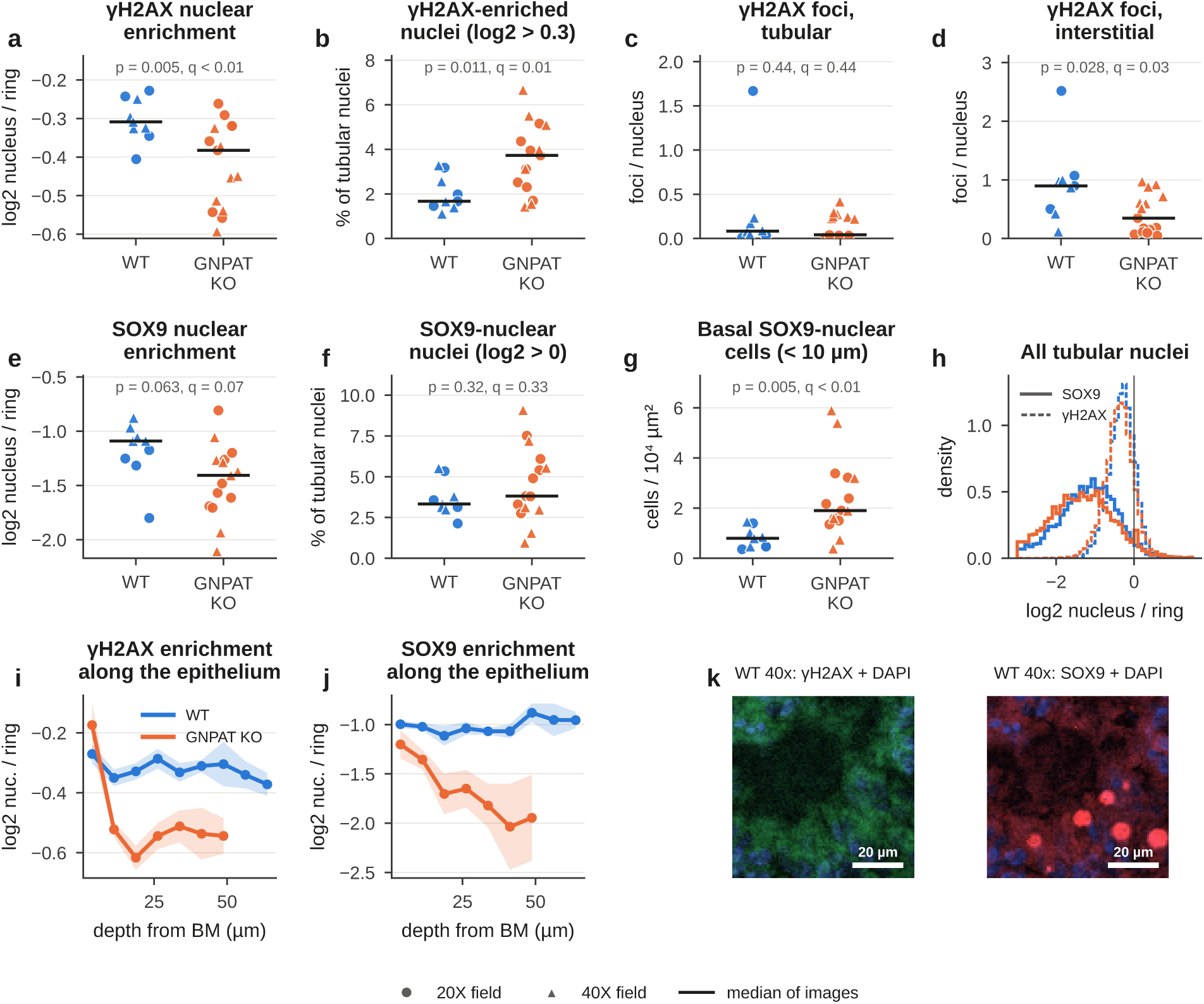
*γ*H2AX and SOX9 readouts. Marker enrichment is *E* = log_2_(nuclear/perinuclear ring intensity), measured within each image; *E* < 0 indicates mainly cytoplasmic signal. **(a)** Median *γ*H2AX enrichment of tubular nuclei per field. **(b)** Proportion of *γ*H2AX-enriched tubular nuclei (*E* > 0.3). **(c, d)** *γ*H2AX foci per tubular (c) and per interstitial (d) nucleus. **(e)** Median SOX9 enrichment of tubular nuclei. **(f)** Proportion of SOX9-nuclear tubular nuclei (*E* > 0). **(g)** SOX9-nuclear cells within 10 µm of the BM per 10^4^ µm² of tubule. In (a–g), symbols are as in Fig. 3, with *p* and *q* (*P*_adj_) printed as in Fig. 3. **(h)** Distributions of SOX9 (solid) and *γ*H2AX (dashed) enrichment for all tubular nuclei, pooled per genotype. **(i, j)** Radial profiles of *γ*H2AX (i) and SOX9 (j) enrichment from the BM towards the lumen in 40× fields (mean ± SEM of fields). **(k)** WT 40× single-marker views: green (*γ*H2AX) with blue (DAPI), left; red (SOX9) with blue (DAPI), right. Canonical meiotic *γ*H2AX patterns and distinct SOX9^+^ basal nuclei are not resolved. Scale bars, 20 µm.

After morphological opening, objects were retained if they were round (equivalent diameter 3.5–12 µm, solidity ≥ 0.85 and eccentricity ≤ 0.85) and isolated, that is, if the mean SOX9-channel intensity in a surrounding 2.5-µm ring was less than 45% of the object’s mean. Each body was assigned to the tubule or interstitium and, if intratubular, given a depth from the BM. These thresholds were set by visual inspection of 40× fields and gave matching densities at 20× and 40×.

For each field, which served as the unit of analysis, 21 metrics were derived from these measurements. They comprised architectural metrics (tubule and interstitium area as a percentage of tissue, and the median inscribed diameter and circularity of valid tubules); epithelial metrics (tubular nuclei per 10^4^ µm² of tubule, median tubular nuclear area, percentage of spermatid-sized nuclei and the 90th percentile of nuclear depth); interstitial metrics (interstitial nuclei per mm² of tissue, median cell territory and percentage of autofluorescent cells in 20× fields); marker metrics (median *γ*H2AX and SOX9 enrichment, the percentages of *γ*H2AX-enriched (*E* > 0.3) and SOX9-nuclear (*E* > 0) tubular nuclei, *γ*H2AX foci per tubular and per interstitial nucleus, and SOX9-nuclear cells within 10 µm of the BM per 10^4^ µm² of tubule); and the density of SOX9-immunoreactive bodies per mm² of tissue, tubule and interstitium. Interstitial nuclei per mm² of interstitium was derived afterwards as the ratio of two of these metrics.

**Figure S6.**
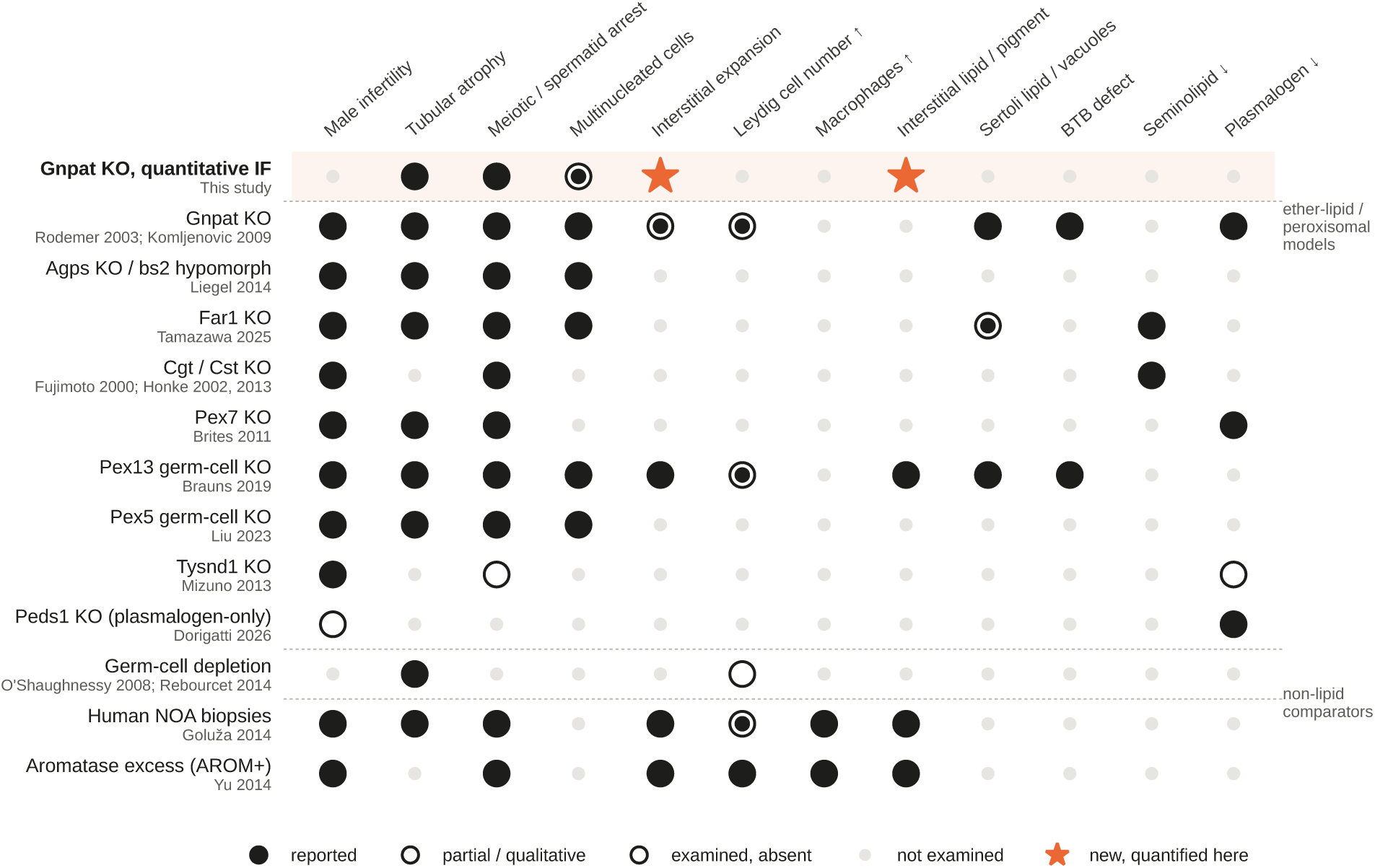
Testis phenotypes of ether-lipid, seminolipid and peroxisomal mutants compared with this study. Each row is a mouse model or human condition, with its source(s) below the model name; each column is a testis feature. Upper block: this study and ether-lipid, seminolipid and peroxisomal models. Lower block: non-lipid comparators (germ-cell depletion by busulfan or diphtheria toxin, human non-obstructive azoospermia, aromatase overexpression). Filled circle, reported; ringed dot, partial or qualitative; open circle, examined and absent; small grey dot, not examined; orange star, new and quantified in this study. Calls are restricted to what each source states (verbatim quotes in Table S2). Komljenovic 2009 contributes the blood–testis-barrier defect of the *Gnpat* row. Abbreviations: gc-KO, germ-cell-specific knockout; NOA, non-obstructive azoospermia; AROM+, aromatase-overexpressing mice; BTB, blood–testis barrier.

### Effect sizes, consistency and sensitivity analyses

For the standardised effect sizes in Fig. 7a, KO − WT differences were divided by the pooled within-magnification standard deviation of WT fields. Consistency between magnifications was assessed by comparing Hedges’ *g* between 20× and 40× fields (Fig. S2d), and threshold sensitivity by recomputing Hedges’ *g* while each analysis threshold was varied over a wide range (Fig. S3). Principal component analysis (scikit-learn) was performed on the 20 metrics available at both magnifications after *z*-scoring each metric across fields. All tests were two-sided.

**Figure S7.**
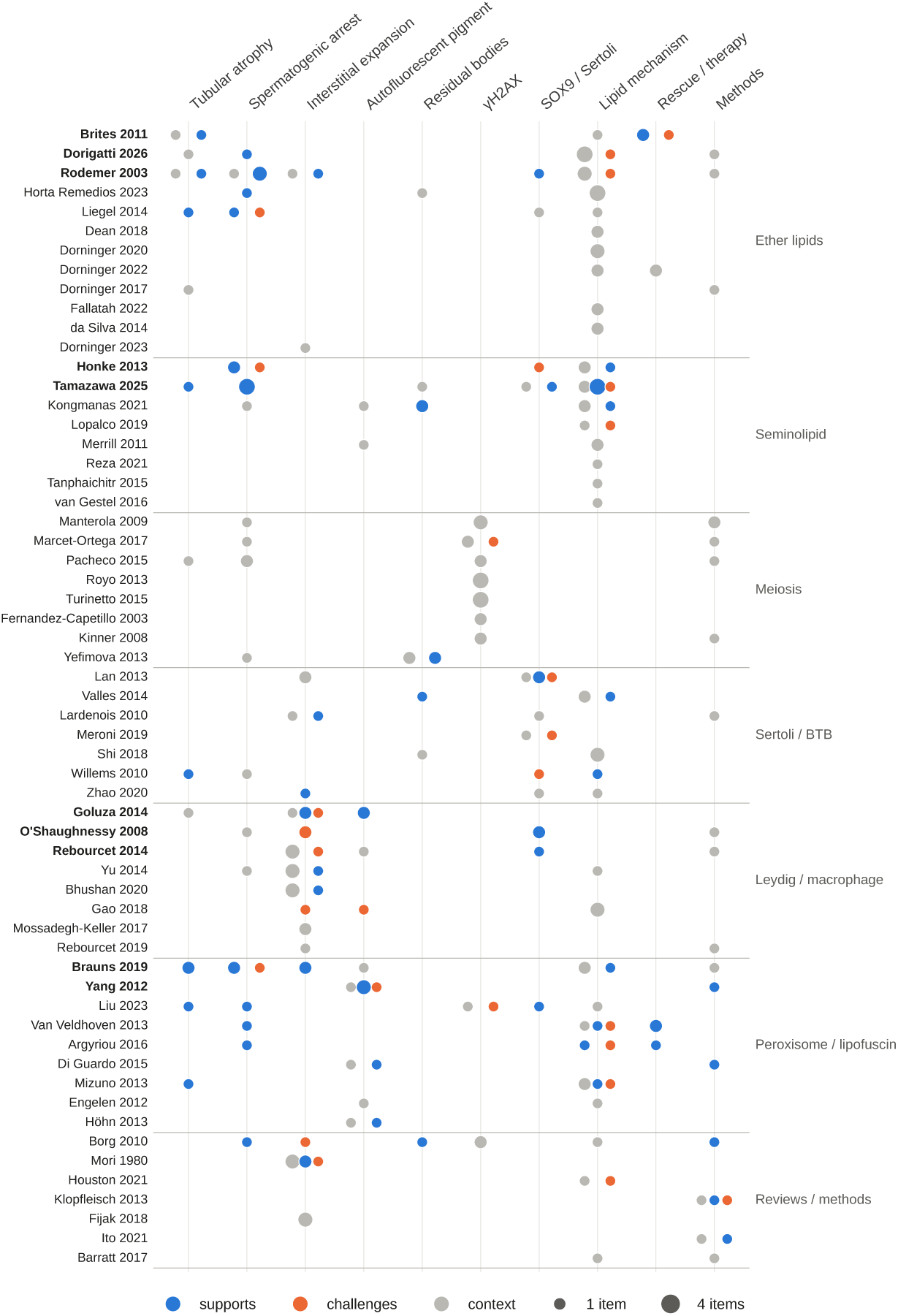
Evidence map of the literature analysed. Each row is one of the 59 papers analysed, grouped by topic (right); papers judged most informative are in bold. Each column is one of our findings. Dots show supporting (blue), challenging (orange) or contextual (grey) evidence items; dot size scales with the number of items. All items, with verbatim quotes and locations, are listed in Table S2.

### Literature synthesis

Open-access primary studies and reviews on ether lipids, seminolipid, meiosis and *γ*H2AX, Sertoli and Leydig cells, testicular macrophages, lipofuscin, peroxisomal disease and male infertility were identified through the Europe PMC REST API (open-access subset, ranked by relevance and citation count). Fifty-eight full texts were retrieved from the PMC open-access collection or from publisher open-access links, and the original description of the *Gnpat* knockout [12] was added. For each paper we recorded the model, the testis phenotype and every statement relevant to our ten findings as a verbatim quote with its location in the source. Every quote was checked programmatically against the full text, bibliographic data were checked against Europe PMC, and each item was classified as supporting, challenging or giving context to a finding. The resulting 265 evidence items are listed in Table S2 and summarised in Figs. S6–S7; phenotype calls in Fig. S6 are restricted to what each source states, and features that were not examined are shown as such.

**Figure S8.**
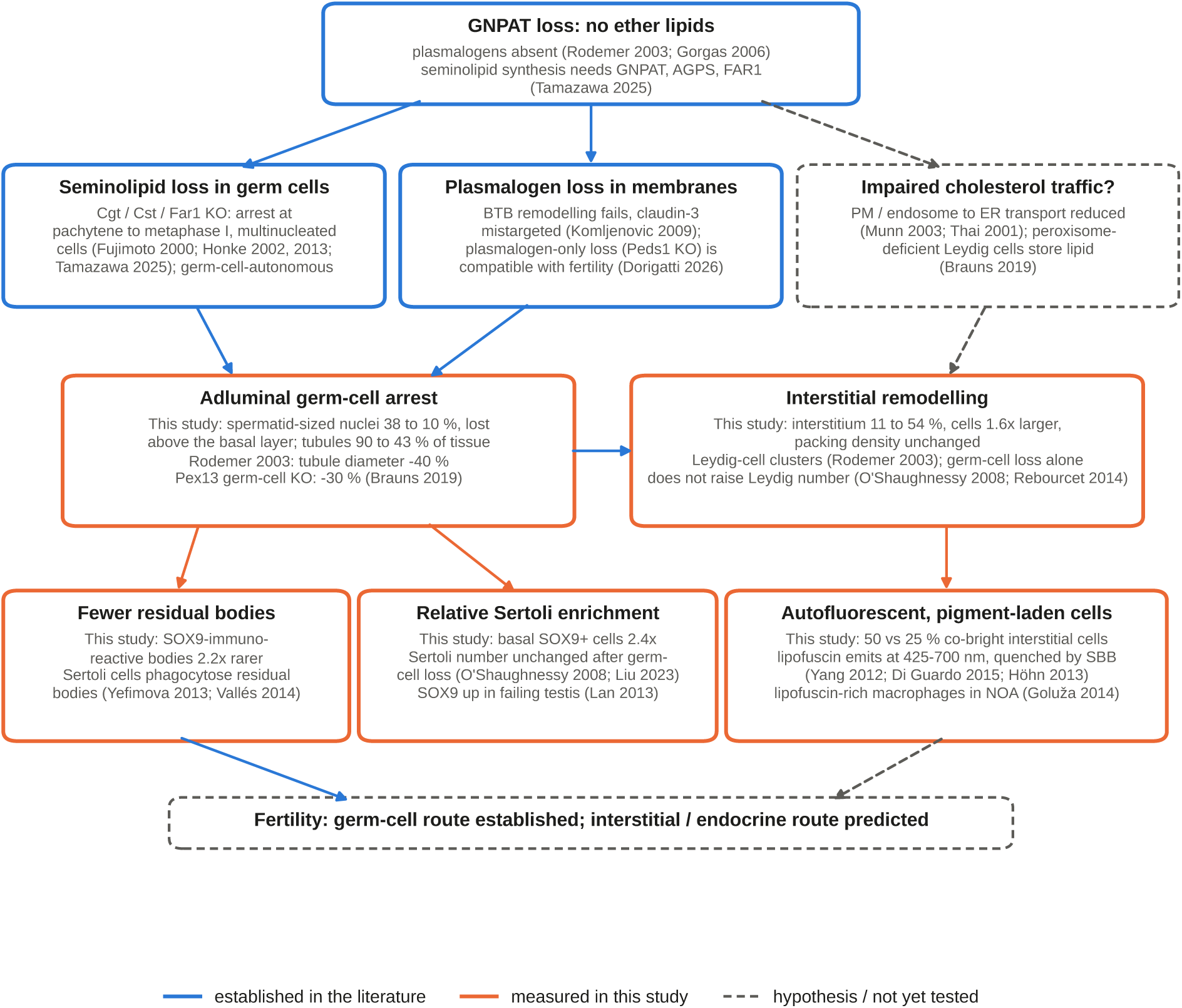
Literature-annotated working model. Boxes summarise the steps from GNPAT loss to the testis phenotype, each with its supporting sources (first author and year; see reference list). Blue boxes and arrows, established in the cited literature; orange, measured in this study; dashed grey, hypotheses not yet tested.

